# H_2_O_2_ repurposes the plant oxygen-sensing machinery to control the transcriptional response to oxidative stress

**DOI:** 10.1101/2024.10.21.619351

**Authors:** Salma Akter, Monica Perri, Mikel Lavilla-Puerta, Beatrice Ferretti, Laura Dalle Carbonare, Vinay Shukla, Yuri Telara, Daai Zhang, Dona M. Gunawardana, William K. Myers, Beatrice Giuntoli, Emily Flashman, Francesco Licausi

**Affiliations:** Department of Chemistry, University of Oxford, Mansfield Road OX1 3TA, Oxford, UK; Department of Biology, University of Oxford, South Parks Road OX13RB, Oxford, UK; University of Bologna, Via Zamboni, 33 40126, Bologna, IT; Department of Biology, University of Pisa, Via Luca Ghini 13, Pisa, IT

**Keywords:** Reactive oxygen species, peroxide, oxidative stress, hypoxia, reoxygenation, desubmergence, plant cysteine oxidases, ethylene response factors

## Abstract

Plants sense reduced oxygen availability (hypoxia) through Plant Cysteine Oxidases (PCOs). Reduced PCO activity in hypoxia, as seen during submergence, stabilises Group VII Ethylene Response Factors (ERFVIIs), master regulators of adaptive metabolic and anatomic responses. Equally important is timely arrest of these responses upon reoxygenation, assumed to occur through ERFVII degradation. Reoxygenation involves reactive oxygen species (ROS) production. Here, we report that instead of degradation, reoxygenation results in ERFVII nuclear stabilisation, an effect mimicked by direct H_2_O_2_ treatment. Interestingly, typical hypoxia marker genes are repressed while genes involved in ROS homeostasis and oxidative stress protection are upregulated. Using *in planta*, heterologous and biochemical assays, we reveal that ROS-related ERFVII stabilisation is caused by PCO inactivation. Stabilised ERFVIIs are retained at hypoxia-responsive promoters but become repressors. Our findings suggest that by responding to both oxygen and ROS, PCOs coordinate ERFVII stability to regulate timely responses to damaging fluctuations in oxygen availability.

## Introduction

All aerobic organisms require oxygen for efficient energy generation; they therefore have mechanisms capable of sensing reduced oxygen availability (hypoxia) to coordinate an adaptive response. Plants experience hypoxia as both an acute stress, such as that caused by submergence, or as a developmental cue in rapidly metabolizing tissues ^1,2^. In both scenarios, the molecular response to the hypoxic conditions has been well characterised and involves reduced activity of oxygen-sensing Plant Cysteine Oxidase enzymes (PCOs) ^3–5^. PCOs create a direct molecular link between environmental O_2_ and stabilization of transcriptional regulators including ERFVIIs ^6^. Using molecular O_2_, PCOs catalyse the oxidation of the N-terminal cysteine residue of their substrates, forming cysteine sulfinic acid (SO_2_H) ^7^. This initiates degradation of the substrate via the Cys/Arg branch of the PRT6 N-degron pathway in normoxic plants ^8^. Alternately, low O_2_ (hypoxia) limits PCO activity resulting in substrate stabilisation ^7^. In the case of ERFVIIs, the stabilized transcription factors bind to hypoxia responsive promoter elements (HRPE) resulting in the elevated expression of a number of core hypoxia responsive genes (HRGs) required to switch from aerobic respiration to fermentative respiration ^9–11^. This adaptive response helps plants survive short periods of hypoxia arising from reduced oxygen diffusion when plants are submerged during a flood event.

While the acute hypoxia experienced by submerged plants is itself stressful, the reoxygenation that occurs on desubmergence is also challenging. Plants must readapt to aerobic respiration and photosynthesis, as well as cope with photoinhibition and desiccation while enduring sugar and energy shortage ^12^. One outcome of hypoxia followed by rapid reoxygenation in both plants and mammals is a burst of elevated reactive oxygen species (ROS), in particular the superoxide anion radical (O_2_^●-^) and hydrogen peroxide (H_2_O_2_) ^13–18^.These ROS are specific molecular derivatives of oxygen produced during normal aerobic metabolism, primarily in chloroplasts and mitochondria with high electron flow and metabolic rates ^19^. Under hypoxia, O_2_^●-^ and H_2_O_2_ production occurs through inefficient O_2_ reduction at electron transport chains, as well as by NADPH oxidases such as RBOHD ^20,21^. Upon reoxygenation, the reactivation of mitochondrial and photosynthetic activities, involving proteins which may have been damaged during hypoxia, causes electron leakage in the electron transport chains and membrane associated processes, which subsequently further increases ROS levels to result in a ROS burst ^22^. There is evidence that this response is associated with hypoxia/reoxygenation in submerged plants: submergence stress in *Arabidopsis thaliana* (hereafter Arabidopsis) is reported to induce systemic ROS wave responses ^23^ and in Arabidopsis rosettes, desubmergence is reported to induce expression of plasma membrane-localized NADPH oxidase/respiratory burst oxidase that produces superoxide in the apoplastic space ^17^.

Given that hypoxia stress entails ROS elevation at its onset and after reoxygenation, there is likely to be cross-talk between cellular responses to both signals. Indeed, in Arabidopsis the hypoxia-responsive universal stress protein 1 (HRU1) is reported to be upregulated by the ROS-related transcription factor heat-shock factor 2A ^24,25^ and ROS-responsive genes are activated by hypoxia-responsive ERFVIIs in Arabidopsis ^26,27^ and SUB1A in rice ^28^. However, it is not known whether there is a direct interaction between ROS and the plant oxygen-sensing machinery. ERFVIIs have been demonstrated to play a crucial role in modulating the response to multiple stresses ^29–31^. It has been proposed that, given the oxygen-sensitivity of the PCOs, reoxygenation would lead to rapid ERFVII degradation and reduced HRG expression ^12^. Here, we investigated the contribution of ROS (specifically H_2_O_2_) ^20^ to PCO-ERFVII mediated responses to hypoxia and reoxygenation. We report that upon reoxygenation or direct H_2_O_2_ treatment, ERFVIIs in fact remain stable and localised in plant nuclei and that this is due to ROS-mediated PCO-inactivation. Interestingly, we find that ERFVIIs repress HRG expression under these conditions while instead activate expression of genes associated with oxidative stress. These data indicate that the PCO-ERFVII signalling pathway can sense both oxygen depletion and a corresponding ROS surge, allowing them to differentiate between hypoxia and oxidative stress and coordinate the appropriate cellular response.

## Results

### ERFVIIs are required for post-hypoxia recovery in Arabidopsis seedlings

Given that plant recovery from a submergence event requires survival of the desubmergence-associated ROS burst, we first investigated whether ERFVII stabilisation during hypoxia contributes to tolerance of this ROS burst and the likely resulting oxidative stress by comparing recovery from reoxygenation in Arabidopsis wild type and *erfVII* mutant plants. We subjected seven-day old seedlings to severe hypoxia (0.5 % O_2_, no light) or normoxia (21 % O_2_, no light) for 24 h; their growth phenotype was analysed immediately after the treatment (before reoxygenation) and 4 days after reoxygenation, during which plants were maintained in 16:8 photoperiod under ambient atmospheric conditions (**Figure 1A, B)**. While we did not observe differences between the wild type and the mutant immediately at the end of the hypoxic treatment, after 4 days of reoxygenation *erfVII* seedlings that had been exposed to hypoxia demonstrated strongly reduced primary root survival in comparison to the wild type (**Figure 1B, C**). Root growth was impaired after reoxygenation in the *erfVII* seedlings (**Figure 1D**), which resulted in reduced biomass accumulation compared to wild type (**Figure 1E**). Although we cannot rule out from this experiment that the differences observed are due to inability of the *erfVII* seedlings to mount an adaptive response to hypoxia, they nevertheless indicated the possibility that ERFVIIs contribute to the plant’s ability to cope with reoxygenation stress.

**Figure 1.**
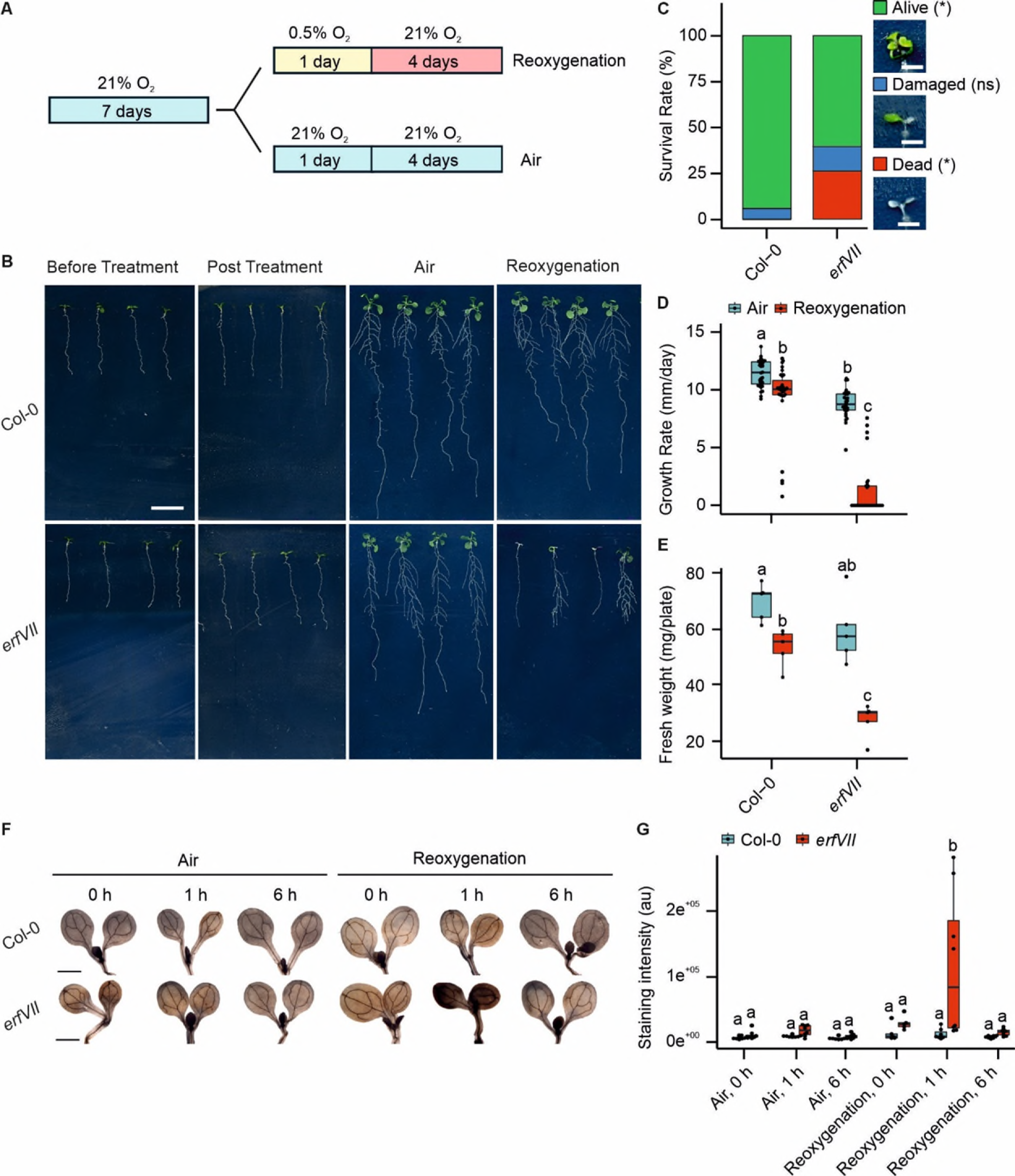
ERFVIIs mediate plant tolerance upon reoxygenation. (**A**) Schematic diagram of experimental design; 7-day old Arabidopsis seedlings were exposed to severe hypoxia (0.5 % O_2_) or air (21 % O_2_), in darkness, for 24 h and subsequently returned to aerobic conditions (16:8 photoperiod) for 4 days to allow for reoxygenation. (**B**) Phenotype of Col-0 and *erfVII* seedlings before and after hypoxia treatment (or air control) and after 4 days reoxygenation (scale bar, 1 cm). (**C**) Percentage of live, damaged or dead seedlings after 4 days of post-hypoxia reoxygenation (*n* = 4, each replicate representing ∼7 seedlings). (**D**) Growth rate of primary roots after 4 days of reoxygenation (*n* = 25-3). (**E**) Fresh weight after 4 days of reoxygenation (*n* = 4, each replicate representing ∼7 seedlings). (**F**) DAB staining of Col-0 and *erfVII* seedlings following exposure to 0.1 % or 21 % O_2_ for 24 h in darkness, and subsequently returned to aerobic conditions (16:8 photoperiod) for 0, 1 or 6 h (scale bar, 0.5 cm). (**G**) Quantification of DAB staining intensity represented in arbitrary unit (au) (*n* = 5-8). Statistical analyses were conducted using: (**C**) the *χ*^2^ test followed by a post-hoc test for survival analysis where asterisks indicate statistical differences between Col-0 and *erfVII*; (**D-G**) two-way ANOVA followed by Tukey HSD test, p<0.05 where different letters indicate statistically different groups.

Based on previous studies where constitutive RAP2.2, RAP2.3 and RAP2.12 protein expression caused increased levels of ROS-related genes in Arabidopsis seedlings ^10^, we speculated that ERFVIIs may function to limit ROS accumulation following reoxygenation. As a marker of ROS accumulation, overall levels of H_2_O_2_ were compared in *erfVII* mutant and wild type seedlings upon reoxygenation, using 3’3’-diaminobenzidine (DAB) staining. Arabidopsis seedlings were treated with 0.5 % O_2_ for 24 h then stained with DAB after 0 h, 1 h and 6 h reoxygenation. Control plants were kept under aerobic conditions for 24 h in the dark, followed by exposure to the light for the same time periods. When compared after reoxygenation, increased DAB staining was observed in *erfVII* seedlings compared to wild type after 1 h, indicating higher accumulation of H_2_O_2_ in *erfVII* than in wild type seedlings upon reoxygenation (**Figure 1F, G**). This result is consistent with a similar study which detected relatively lower levels of H_2_O_2_ upon desubmergence in the aerial tissue of rice expressing *SUB1A*, an N-degron pathway resistant ERFVII, compared with plants lacking *SUB1A* ^28^. Together, the results suggest that ERFVIIs contribute to recovery post-submergence by reducing accumulation of ROS upon reoxygenation.

### ERFVIIs are stabilised in Arabidopsis seedlings by both post-hypoxia reoxygenation and oxidative stress

During hypoxic conditions, ERFVIIs are known to be stabilized and accumulate in the nucleus ^32,33^ (**Figure 2A**). Since our results indicate that ERFVIIs also contribute to growth recovery after reoxygenation, we investigated the abundance and localization of RAP2.12 in Arabidopsis cells in conditions that mimic post-submergence reoxygenation, as well as upon the addition of exogenous ROS in the form of tert-butyl hydroperoxide (TBHP) ^34,35^. We treated transgenic Arabidopsis seedlings constitutively expressing RAP2.12 fused with green fluorescent protein (GFP) with or without TBHP (1 mM) in normoxia, hypoxia and after reoxygenation (3 h post-hypoxia, consistent with the timing of oxidative damage of lipids reported in Arabidopsis after desubmergence ^17^). We then looked for RAP2.12-associated GFP signal in Arabidopsis root cells using confocal imaging. As expected, we observed elevated RAP2.12-GFP localization in the nuclei of hypoxic plants compared to normoxic plants (**Figure 2B**). Upon reoxygenation, ERFVII may be presumed to destabilize with the resumption of PCO enzyme function in aerobic conditions, however we found that the RAP2.12-GFP signal persisted in the nucleus after 3 h reoxygenation treatment, both in the presence and absence of TBHP. Interestingly, we observed that TBHP treatment resulted in an increase in RAP2.12-GFP signal in the nuclei of plants exposed to both air and hypoxia (**Figure 2B**). We confirmed these observations using a second transgenic Arabidopsis line expressing RAP2.3 fused with GFP (**Figure S1A**). A time-course observation of TBHP-treated, normoxic plants showed stabilisation of RAP2.3 within less than 2 h (**Figure S1B**). Collectively, these data revealed that ERFVIIs remain stable in the nucleus upon reoxygenation, when plants are exposed to ROS as well as O_2_.

**Figure 2.**
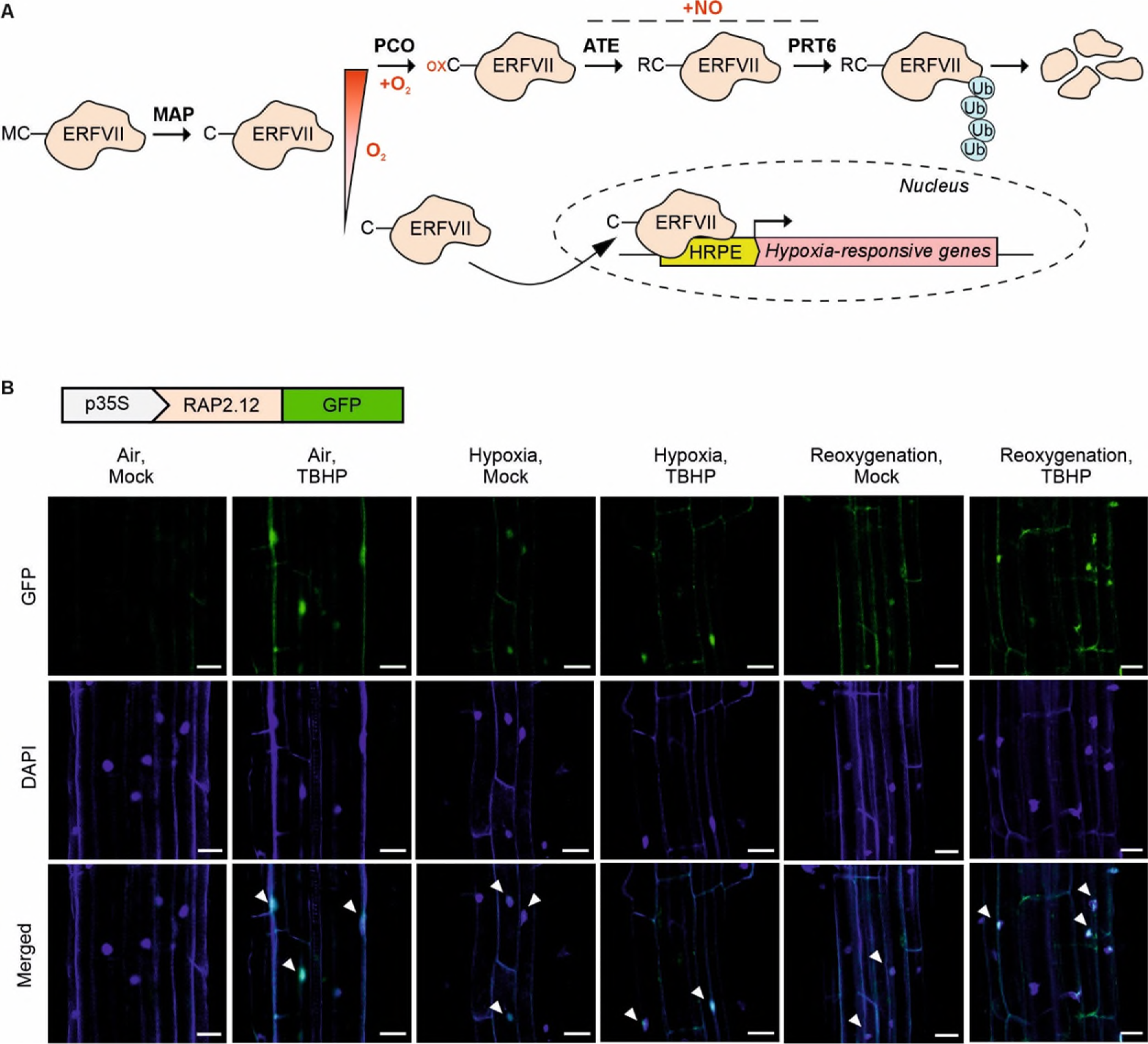
RAP2.12 localizes to the nucleus upon oxidative stress as well as during reoxygenation. (**A**) Schematic representation of N-degron pathway-controlled destabilisation of ERFVII. Methionine (M) in the N-terminal position is target of the methionine aminopeptidase enzyme (MAP), leaving an exposed cysteine which, in presence of oxygen (O_2_) is targeted for oxidation by plant cysteine oxidase (PCO). The resulting Cys-sulfinic acid is sequentially modified by arginyl aminotransferases 1/2 (ATE) triggering proteasomal degradation after ubiquitination by proteolysis E3 ligase (PRT6), in presence of nitric oxide (NO). Upon hypoxic conditions, ERFVII are stabilized and relocalize to the nucleus where they bind the hypoxia responsive promoter element (HRPE) triggering the hypoxic response. Additional abbreviation: M, methionine; C, cysteine; oxC, oxidized cysteine; R, arginine; Ub, ubiquitin. (**B**) Localization of 35S:RAP2.12-GFP (green) in 7-day old Arabidopsis seedlings upon 1 mM TBHP treatment in normoxia (21% O_2_ v/v O_2_/N_2_), hypoxia (1% O_2_ v/v O_2_/N_2_) or after 3 h of reoxygenation. Nuclear localization is confirmed using DAPI staining (blue). White arrows indicate nuclear colocalization (scale bar, 50 µM).

To complement the data obtained with GFP-fused ERFVIIs, we generated an Arabidopsis line constitutively expressing the full RAP2.3-coding sequence fused in frame with the nano-luciferase enzyme (RAP2.3-nLUC). This transgenic line conveniently reports on ERFVII stability. Seven-day old RAP2.3-nLuc seedlings were treated with or without TBHP (1 mM) in normoxia, hypoxia and after reoxygenation, as in the subcellular localisation experiment. As expected, in the absence of TBHP luciferase activity was higher in hypoxic seedlings than in normoxic ones, consistent with N-degron pathway-mediated degradation of RAP2.3 in normoxia (**Figure 3A**). TBHP-treated seedlings in normoxia exhibited significantly higher luciferase activity compared to controls, consistent with ERFVII stabilisation under aerobic conditions when cells are exposed to H_2_O_2_ (**Figures 2B, 3A**). We confirmed that this phenomenon is dose- and time-dependent (**Figure 3B, C**) with RAP2.3 stabilization reaching a plateau at 2 h of treatment **(Figure 3C)**. Furthermore, during reoxygenation, and in the absence of exogenous TBHP, luciferase activity did not decrease from hypoxic levels, remaining significantly higher than in normoxic seedlings (**Figure 3A**). Taken together, these measurements quantitatively confirm the stability of ERFVIIs upon both oxidative stress and during the reoxygenation period. A second ERFVII reporter line expressing a 1-28 aa fragment of RAP2.12 fused with the firefly luciferase enzyme (RAP2.12_1-28_-FLuc)^3^ exhibited similar behaviour (**Figure S2A**), with elevated FLuc activity in TBHP-treated aerobic and reoxygenated seedlings, further confirming our findings.

**Figure 3.**
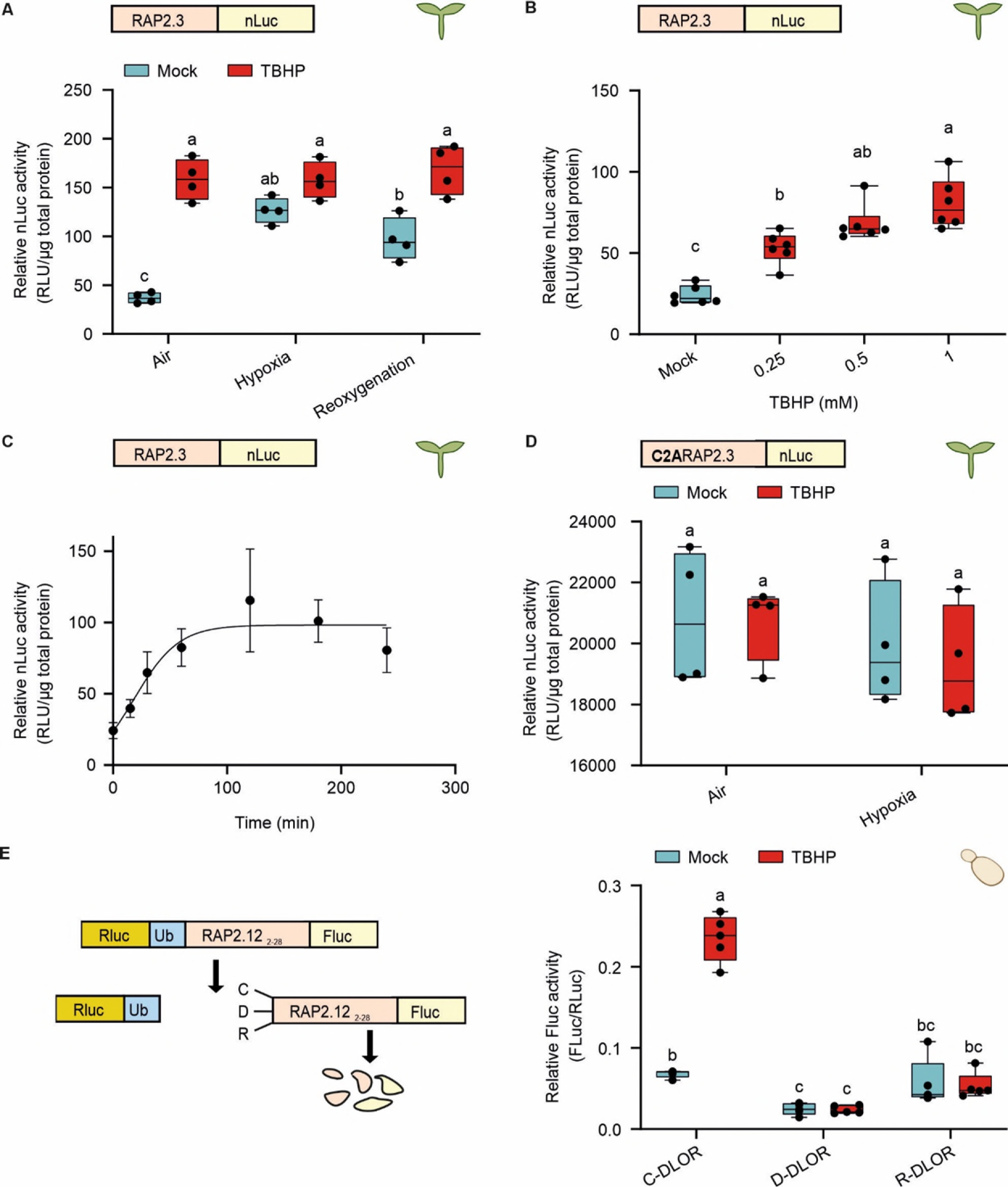
Oxidative stress increases ERFVII stability in a N-degron pathway-dependent manner. (**A**) Relative nLuc activity of 35S:RAP2.3-nLuc seedlings treated under hypoxic (1% O_2_ v/v O_2_/N_2_) or aerobic (21 % O_2_ v/v O_2_/N_2_) conditions for 6 h, followed by 3 h reoxygenation, 1 mM TBHP or mock treatment (*n* = 4). (**B**) Relative nLuc activity of 35S:RAP2.3-nLuc seedlings treated with different TBHP concentrations upon aerobic conditions over 4 h (*n* = 6). (**C**) Relative nLuc activity of 35S:RAP2.3-nLuc seedlings treated with 1 mM TBHP upon aerobic conditions measured over 6 h (*n* = 6). (**D**) Relative nLuc activity of 35S:C2ARAP2.3-nLuc variant upon 1mM TBHP or mock treatment, under hypoxic or (1 % O_2_ v/v O_2_/N_2_) or aerobic (21 % O_2_ v/v O_2_/N_2_) conditions (*n* = 4). (**E**) Relative FLuc activity in yeast cultures expressing *At*PCO4 together with either C-DLOR, R-DLOR or D-DLOR and exposed to 0.5 mM TBHP or mock treatment for 30 min. Statistical differences were evaluated using One-way ANOVA (**B**) or Two-way ANOVA (**A, D, E**) followed by Tukey HSD test (p < 0.05).

### ROS-induced ERFVII stabilisation is conferred via the N-terminus

The observations obtained with the RAP2.12_1-28_-FLuc line indicate that the prevention of the predicted O_2_-dependent degradation exclusively required the N-terminal region of ERFVIIs, known to contain the conserved N-degron MCGGAII ^32^. We therefore sought to determine whether H_2_O_2_-mediated ERFVII stabilisation is conferred via modification of its Nt-Cys (the catalytic target of the PCOs) rather than other ERFVII residues. We used a transgenic RAP2.3-nanoluciferase fusion in which the N-terminal cysteine of the transgenic RAP2.3 is mutated to alanine (C2ARAP2.3-nLUC), thereby preventing PCO-initiated proteolysis of RAP2.3. We examined the luciferase signal in C2ARAP2.3-nLUC seedlings exposed to air or hypoxia and treated with or without TBHP. In contrast to Cys2-RAP2.3-nLuc (**Figure 3A-C**), the luciferase signal of C2A-RAP2.3nLUC did not vary significantly upon hypoxia or TBHP treatment (**Figure 3D**). This suggested that prevention of ERFVII degradation is specifically due to the inability of the N-degron pathway to process protein N-termini and that TBHP-induced oxidative stress cannot increase ERFVII stability when ERFVIIs are no longer N-degron substrates.

The comparative observations of luciferase activity in RAP2.3-, RAP2.12_1-28_- and C2A-RAP2.3-linked reporter lines suggested that ERFVII stabilization upon TBHP treatment or upon post-hypoxia reoxygenation is linked to the protection of N-terminal cysteine from the N-degron pathway. To dissect the step at which N-degron processing is inhibited under these conditions, we next used a heterologous DLOR (Double Luciferase Oxygen Reporter) reporter assay developed in the budding yeast *Saccharomyces cerevisiae* ^36^. Here, the relative luciferase output (FLuc/RLuc) of the DLOR reporter is strictly dependent on transgenic PCO enzymes (in this case *At*PCO4) expressed in yeast, which otherwise lack N-terminal cysteine oxidation activity. TBHP treatment of exponentially grown yeast caused a fast and dose-dependent increase of DLOR output, thus demonstrating TBHP-induced ERFVII stabilization, similar to our observations in plants (**Figures 3E, S2B, C**). The optical density (OD) of the TBHP-treated cultures was reduced slightly in TBHP-treated cells compared to control, but this was dose-independent in contrast to the dose-dependent effect on FLuc activity, indicating a specific effect of TBHP on the DLOR system (**Figure S2D**).

Having confirmed the stabilisation of ERFVII in PCO-expressing yeast, we then used this system to independently probe each step of the N-degron pathway for susceptibility to TBHP. The Arg/N-degron pathway in yeast is mediated by ATE1-dependent arginylation of Nt-Glu-, Nt-Asp- or Nt-Cys-SOO(O)H-initiating proteins and UBR1-catalyzed ubiquitinylation of Nt-Arg-proteins ^8,37^. We used two alternative versions of DLOR, D-DLOR and R-DLOR, in which the Nt-Cys (C) of the RAP2.12_2-28_ domain is mutated to aspartate (D) or arginine (R), respectively ^38^. C-DLOR, D-DLOR and R-DLOR were all treated with 0.5 mM TBHP for 30 min and the relative FLuc activity was measured (**Figure 3E**). The signal was increased by TBHP treatment only in C-DLOR yeast, whereas it did not change significantly in D-DLOR and R-DLOR strains (**Figure 3E**). These results indicate that TBHP does not interfere with ATE1 and UBR1 function, thereby suggesting that ROS-induced ERFVII stabilization in yeast occurs through inhibition of the PCO-catalysed step, suggesting that this is also the case in plants.

### H_2_O_2_ inhibits recombinant PCO enzymes

We set out to determine the biochemical mechanism that leads to inhibition of Nt-Cys sulfinylation in plant cells. We first examined whether TBHP or H_2_O_2_ treatment could inhibit PCO activity. We treated 10 µM recombinant AtPCO4 with 50 µM TBHP or H_2_O_2_ and then tested its ability to catalyse oxidation of a 14-mer peptide corresponding to the RAP2.12 N-terminus (hereafter RAP2_2-15_) using liquid-chromatography mass spectrometry (LC-MS) as described previously ^38^. Both treatments led to a significant reduction in AtPCO4-catalysed RAP2_2-15_ oxidation compared to untreated enzyme (**Figure 4A**). In a dose-dependence assay, we found 3 to 5 μM H_2_O_2_ to be sufficient to inhibit 50 % activity of 10 μM AtPCO4 (**Figure 4B**). We also assessed the sensitivity of all five AtPCOs to H_2_O_2_, in a 5-minute end point assay with 5 µM or 10 µM H_2_O_2_ (**Figure 4C**) and found that all of these enzymes showed significantly decreased activity in response to H_2_O_2_ exposure, with strongest inhibition for AtPCO4 and AtPCO5. Given that Cys residues are susceptible to non-enzymatic oxidation, we also tested whether H_2_O_2_ could directly oxidise RAP2_2-15_ under the same conditions and thus generate potential N-degrons. We found that 1mM H_2_O_2_ treatment resulted in a small amount of spontaneous RAP2_2-15_ Nt-Cys oxidation (**Figure 4A**), comprising sulfenic acid (+16 Da), sulfinic acid (+32 Da) or sulfonic acid (+48 Da) (**Figure S3**). Under these conditions, direct H_2_O_2_ treatment only resulted in 5.3 % RAP2_2-15_ oxidation compared to AtPCO4-catalysed oxidation. Overall, therefore, the results indicate that the primary effect of H_2_O_2_ and TBHP on N-degron pathway mediated ERFVII stability is inhibition of PCO activity.

**Figure 4.**
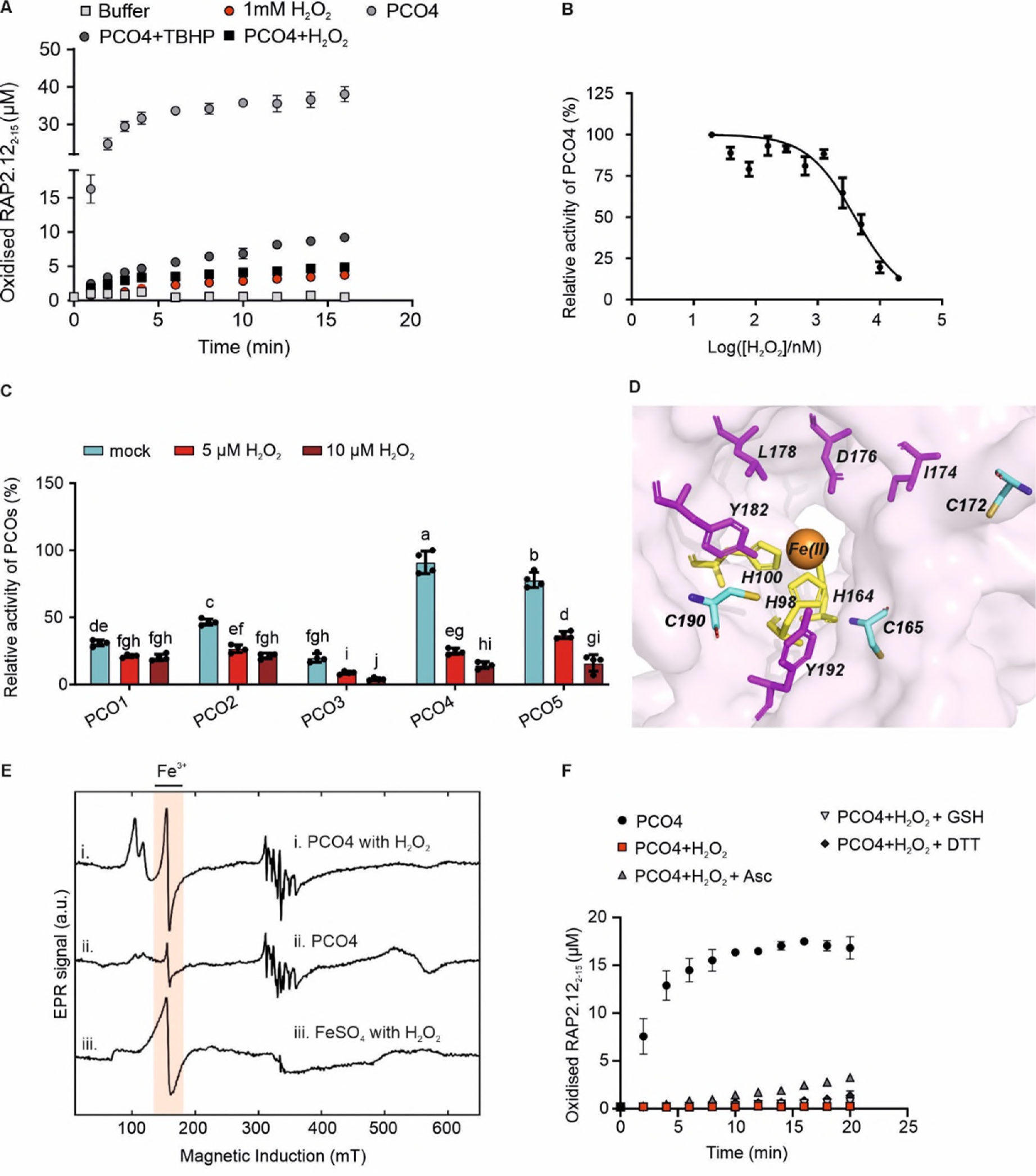
H_2_O_2_ inhibits recombinant PCO enzymes. (**A**) The activity of 50 µM H_2_O_2_- and 50 µM TBHP-treated or non-treated recombinant PCO4 enzyme-catalysed oxidation of RAP2_2-15_. Non-enzymatic RAP2_2-15_ oxidation by 1 mM H_2_O_2_ was included to compare with the oxidation of RAP2_2-15_ by PCO4. (**B**) H_2_O_2_ dose-dependent effect on PCO4 enzyme activity. (**C**) H_2_O_2-_mediated inhibition of PCOs1-5; statistical differences were evaluated using Two-way ANOVA followed by Tukey HSD test (p < 0.05). Error bars display standard deviation (S.D.) (*n* = 3). (**D**) Active site view of crystal structure of AtPCO4 (PDB: 6S7E); Fe cofactor (orange sphere) is bound by a triad of His residues His98, His100, and His164 (highlighted in yellow), Cys residues found to be oxidised by H_2_O_2_ shown in cyan. (**E**) X-band CW-EPR spectra of (i) AtPCO4 with H_2_O_2_, (ii) AtPCO4 only and (iii) FeSO_4(aqueous)_ with H_2_O_2_. Spectrum in (i) shows Fe(III) signal intensity at 150 mT similar to that seen in (iii), but with signal splitting at g_eff_ = 4.29, g′_eff_=6.4 and g″_eff_=5.7, first as typical rhombic coordination of Fe(III) with two axially-symmetric species. (**F**) Activity restoration test of H_2_O_2_ treated PCO4 enzyme, using known cellular reductants glutathione (GSH), ascorbate (Asc) and dithiothreitol (DTT).

To explore the mechanism of H_2_O_2_-mediated PCO enzyme inhibition, we first tested whether H_2_O_2_ causes oxidative modification of the PCO4 enzyme. Whole protein mass spectrometry of H_2_O_2_-treated enzyme revealed a +209 Da mass increase (**Figure S4A)** which we considered likely due to non-specific oxidation of susceptible residues. Cys residues are particularly prone to oxidative modification to Cys-sulfinic acid, therefore we used a sulfinic acid-specific DiaAlk probe ^39^ which confirmed that Cys residues of PCO4 undergo sulfinic acid modification upon H_2_O_2_ treatment (**Figure S4B)**. Further interrogation using LC-MS/MS based proteomic analysis of H_2_O_2_ treated and non-treated PCO4 showed that Cys165, Cys172 and Cys190 of PCO4 form sulfinic (+31.98 Da) and sulfonic acids (+47.97), all of which are close to the AtPCO4 active site (**Figure 4D**), with the highest proportion of oxidation occurring on Cys165 upon H_2_O_2_ treatment (**Figure S4C-F)**. Oxidation of any of these Cys residues could therefore potentially negatively impact the ability of PCO4 to bind substrate or catalyse oxidation. The PCOs are Fe(II)-dependent thiol dioxygenases ^7^ (**Figure 4D**), therefore as well as protein modification, ROS-mediated oxidation of PCO active site Fe(II) could result in their inhibition. Continuous wave electron paramagnetic resonance (CW-EPR) spectroscopy revealed that H_2_O_2_ treatment resulted in an increase in rhombic Fe(III) signal at the PCO4 active site, similar to that seen upon treatment of isolated Fe(II)._aqueous_ with H_2_O_2_ (**Figure 4E**). This confirmed that Fe(II) oxidation is indeed another possible mechanism for PCO inhibition. Finally, we tested whether the H_2_O_2_ mediated inhibition of AtPCO4 could be restored using known cellular reductants glutathione and ascorbate ^40,41^ as well as a synthetic thiol reductant dithiothreitol ^42^. None of the reductants were able to recover the activity of H_2_O_2_-treated enzyme (**Figure 4F**), consistent with the duration of ERFVII stabilisation seen *in vivo*.

Collectively, the results indicate non-specific impacts of H_2_O_2_ treatment on PCO function, likely combining both catalytic inactivation and impacts on protein structure, which together result in the ERFVII stabilisation observed *in vivo*.

### ROS-triggered oxidative stress disrupts hypoxia-responsive gene expression

Our results indicate that elevated ROS, as occurs in reoxygenation, causes inhibition of PCO activity, leading to stabilization of ERFVII. This unexpected result contrasts with previous reports that show rapid repression of hypoxia-inducible genes upon reoxygenation ^43,44^ as well as lack of activation of the same set of genes under oxidative stress conditions ^45^. To address this conundrum, we set out to investigate whether ROS signalling affects the hypoxia response in Arabidopsis by measuring the expression of hypoxia-responsive genes (HRGs) upon hypoxia, reoxygenation and direct treatment with TBHP. We exposed 7-day old seedlings to 1 mM TBHP or mock control under normoxia, hypoxia and reoxygenation. We monitored the expression of six hypoxia marker genes: three involved in fermentative metabolism (*Alcohol Dehydrogenase 1* (*ADH1*), *Pyruvate Decarboxylase 1* (*PDC1*) and *sucrose synthase 1 (SUS1*)), the *non-symbiotic hemoglobin 1 (Hb1)* and two transcription factors (*HYPOXIA RESPONSE ATTENUATOR1 (HRA1),* and *LATERAL ORGAN BOUNDARY DOMAIN 41 (LBD41))* ^11^, using real-time qPCR. This analysis confirmed upregulation of all six genes in hypoxia and their downregulation under reoxygenation conditions (**Figure 5A**), consistent with findings from previous studies ^43^. Interestingly, TBHP-treated plants showed significantly lower expression levels of all hypoxia responsive genes compared to untreated plants under hypoxia (**Figure 5A**), with expression levels similar to aerobic conditions. These genes also showed low expression levels in reoxygenated plants treated with TBHP. These results are surprising given that we observed ERFVII stabilisation on TBHP treatment and/or reoxygenation and suggested that ROS interfere with the plant’s hypoxia responses, even while plants are experiencing hypoxia stress. Consistent with these observations, expression of the core hypoxia responsive genes in the *RAP2.3-nLUC* transgenic Arabidopsis lines was also elevated in hypoxia, but significantly decreased upon TBHP-treatment under hypoxia and/or under reoxygenation (**Figure S5**). To determine whether TBHP-treatment promoted expression of oxidative stress responsive genes, we investigated expression of Cysteine-Rich Receptor-Like Kinase (*CRK36*), GLUTATHIONE S-TRANSFERASE (CLASS TAU) 24 (*GSTU24*) and zinc-finger protein (*ZAT12*) in Arabidopsis wild type 7-day old seedlings under similar treatments (**Figure 5B)**. These genes are reported to be triggered by ERFVII transcription factors ^26^. We found that all three genes were upregulated upon TBHP treatment (**Figure 5B**). These results confirmed that the plants did indeed experience oxidative stress as a result of TBHP treatment, and demonstrate that TBHP treatment does not inhibit their transcriptional machinery, but selectively acts to prevent expression of HRGs.

**Figure 5.**
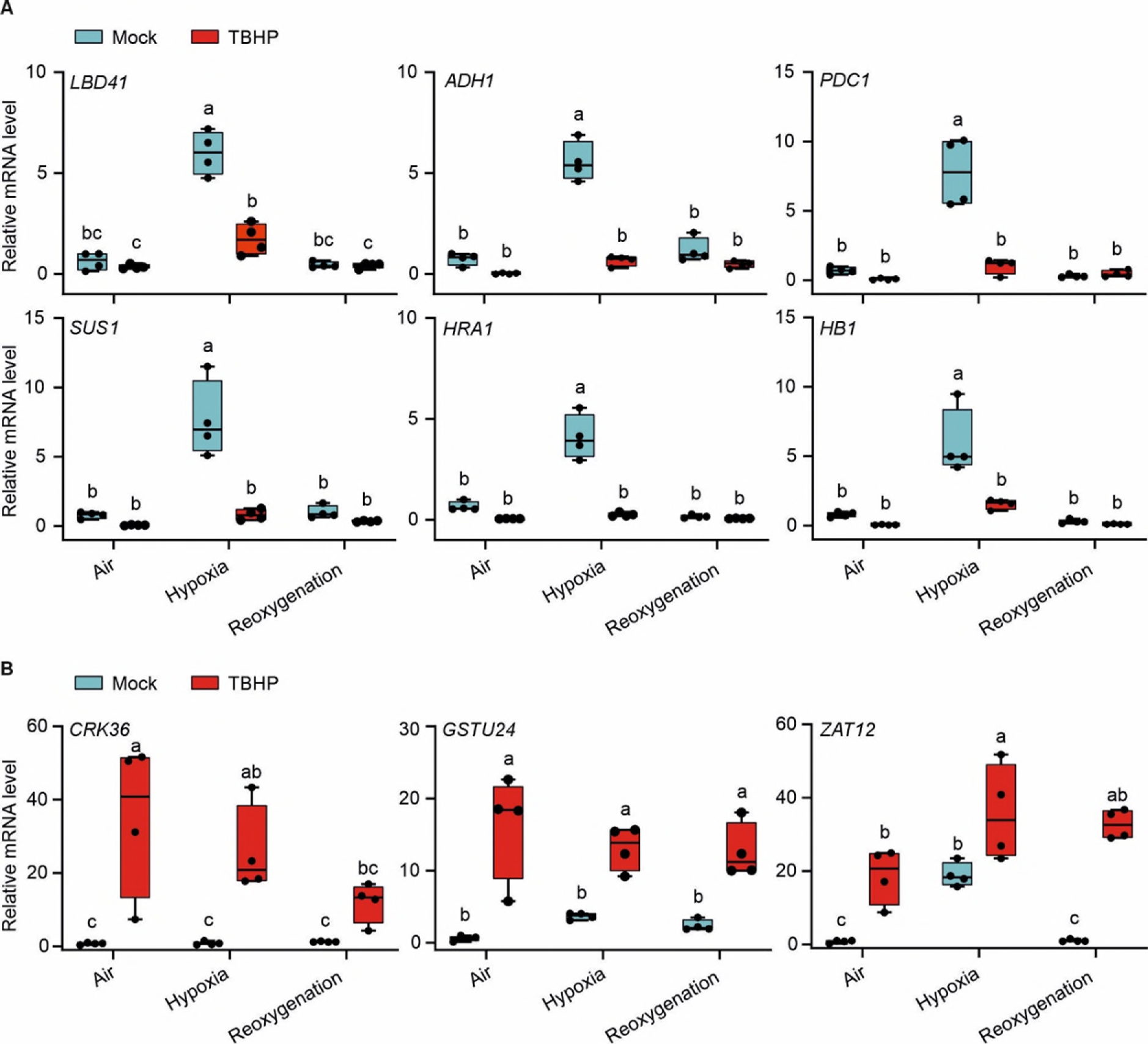
ROS-triggered oxidative stress disrupts hypoxia-responsive gene expression, without damaging the transcriptional machinery. (**A**) Relative expression level of six hypoxia-responsive genes *LBD41*, *ADH1*, *PDC1*, *SUS1*, *HRA1*, *HB1* in air, hypoxia or reoxygenation, upon 1 mM TBHP or mock treatment in *Col-0* seedlings (*n* = 4). (**B**) Relative expression level of ROS-responsive genes *CRK36*, *GSTU24* and *ZAT12* in in air, hypoxia or reoxygenation, upon 1mM TBHP or mock treatment (*n* = 4). Statistical analyses were conducted using Two-way ANOVA followed by Tukey HSD test, p<0.05. Different letters indicate statistically different groups.

HRGs are controlled by ERFVIIs through their interaction with the Hypoxia Responsive Promoter Element (HRPE) ^10,46,47^. To understand how TBHP treatment resulted in reduction of HRG expression, we investigated whether oxidative stress reduces the binding of ERFVIIs to HRPEs. To do this, we used a transgenic Arabidopsis line with a synthetic promoter harbouring a five time repeat of HRPE driving the expression of a gene encoding a nanoluciferase enzyme (*nLuc*). We measured the expression level of *nanoluciferase* as well as the expression of the same six core hypoxic genes described above, in air, hypoxia and reoxygenation with or without TBHP-mediated oxidative stress (**Figure 6A, Figure S6**). We found that *nLuc* expression was upregulated under hypoxic conditions, but downregulated upon oxidative stress, similar to the HRGs (**Figure 6A**, **Figure S6**). These results indicate that TBHP treatment interferes with the transactivation of genes whose promoter includes at least one HRPE, and not through a different DNA motif.

**Figure 6.**
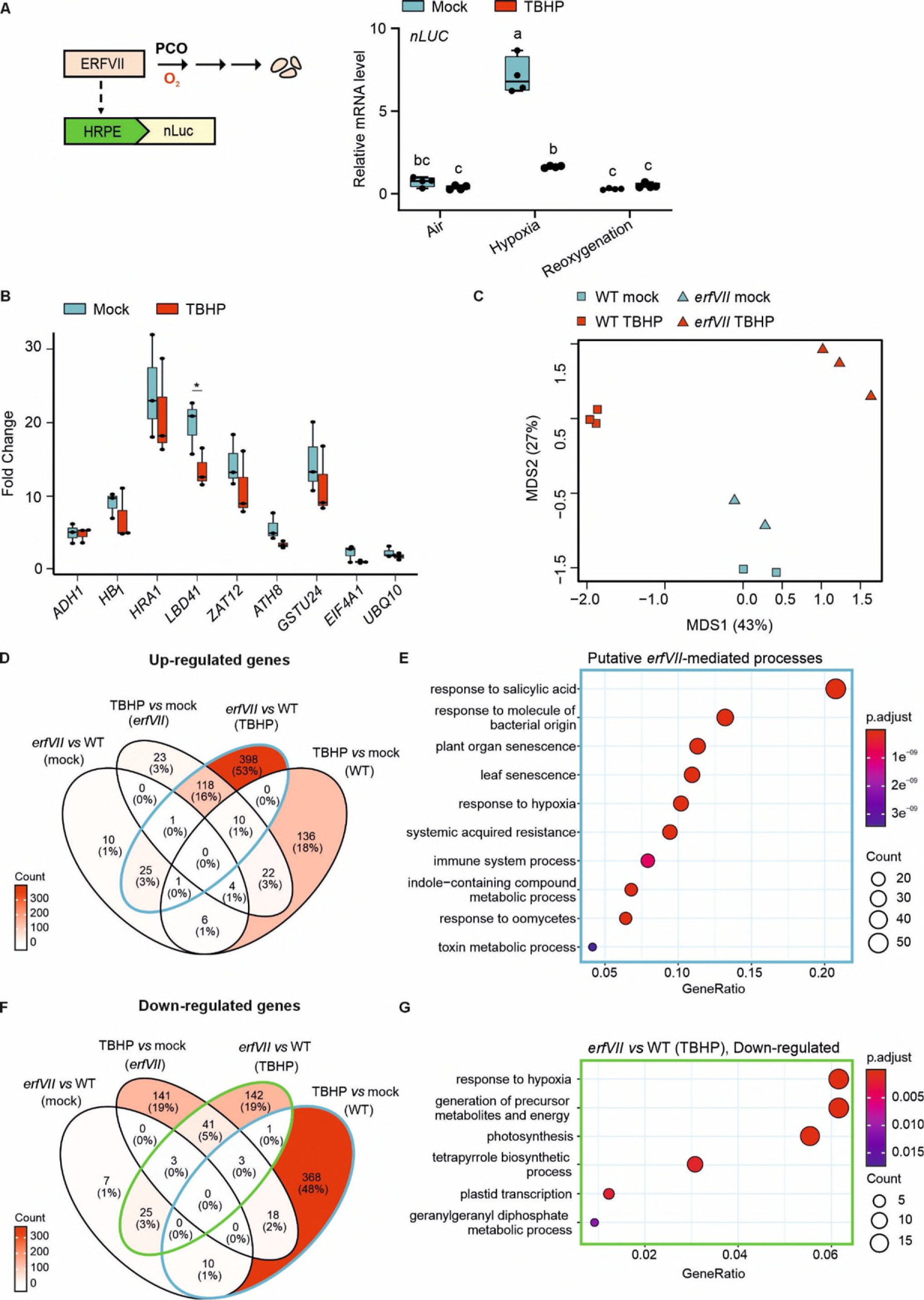
Oxidative stress does not affect the ERFVII DNA binding affinity and induces an ERFVII-dependent transcriptome change. (**A**) Relative expression of *nLuc* in HRPE:nLuc seedlings treated with 1 mM TBHP or mock under air, hypoxia or reoxygenation (*n* = 4). Statistical analyses were conducted using Two-way ANOVA followed by Tukey HSD test, p<0.05. Different letters indicate statistically different groups. (**B**) ChIP-qPCR analysis of Δ13RAP2.12-GFP for hypoxia-responsive genes (*ADH1*, *HB1*, *HRA1*, *LBD41*) and oxidative stress genes (*ZAT12*, ATH8*, GSTU24*) promoters, compared to negative controls (*EIF4A1*, *UBQ1O*). Asterisks indicate statistically significant difference between the two mean values for each gene (Student’s t test, p < 0.05, *n*=3). (**C**) Multidimensional scaling plot (MDS) conducted on the normalized gene expression values of *erfVII* and Col-0 seedlings treated with TBHP or mock. Horizontal and vertical coordinates show MDS1 and MDS2, respectively, with the amount of variance contained in each component (43% and 27%). Each point in the plot represents a biological replicate. Differences in symbols and colour indicate difference in genotype and treatment, respectively. (**D**) Venn Diagram indicating the overlap of up-regulated genes for *erfVII* or Col-0 treated upon TBHP or mock conditions. (**E**) GO enrichment results for biological process of genes that are up-regulated genes in *erfVII* seedlings compared to wild-type, upon TBHP treatment, and down-regulated in the wild-type upon TBHP treatment compared to mock. (**F**) Venn Diagram indicating the overlap of down-regulated genes for *erfVII* or Col-0 treated upon TBHP or mock conditions. (**G**) GO enrichment results for biological process of the down-regulated genes in TBHP-treated *erfVII* seedling compared to wild-type. Circle size indicates the gene count per GO term, with color maps indicating the False Discovery Rate (FDR) value (p.adjust).

We next raised the question of whether oxidative stress hinders the ability of ERFVII to physically interact with the HRPE. We performed a chromatin immunoprecipitation (ChIP) experiment using a GFP-tagged truncated version of RAP2.12 lacking 13 amino acid residues at the N-terminus (Δ13RAP2.12-GFP, ^26,32^). Due to the absence of the Cys-residue at the N-terminus, RAP2.12 is constitutively stable and therefore able to transactivate the hypoxia response. 7-day old seedlings were treated with TBHP upon normoxic conditions, followed by evaluation of the enrichment in the promoter regions of four hypoxia-responsive genes, *ADH1*, *HB1*, *HRA1* and *LBD41*, together with three ERFVII-dependent oxidative stress-responsive genes *ZAT12*, *ATH8* and *GSTU24* (**Figure 6B**). *UBQ10* and *EI4A1* promoters, which do not respond to ERFVII, were used as negative controls. ChIP-qPCR revealed an enrichment for all genes upon normoxic conditions, when compared to the negative controls, confirming that Δ13RAP2.12-GFP binds to the promoter regions of these genes. A significant reduction in genomic enrichment was only observed for the *LBD41* promoter upon TBHP treatment, while no statistical difference was measured for any other promoters. These results suggest that oxidative stress only mildly affects RAP2.12 binding to selected genomic regions. Additional regulation is therefore likely required to reach the complete suppression of the transcriptional hypoxic response observed in this study (**Figure 6B**, **Figure 5A**).

### ERFVIIs control transcriptional reprogramming during oxidative stress

Having shown that ERFVII stability upon oxidative stress does not induce the canonical hypoxia response, we set out to investigate more broadly the contribution of ERFVII transcription factors to the transcriptional response to oxidative stress. To this end, we sequenced the polyA-enriched transcriptome of 7-day old wild type and *erfVII* seedlings treated with 1mM TBHP. We included untreated seedlings of both genotypes as controls.

Multidimensional scaling (MDS) analysis (**Figure 6C**) confirmed a moderate effect of ERFVII inactivation under control conditions ^48^, where only 92 genes were differentially expressed (|FC|>1.5, p<0.05) between the mutant and the wild type (**Fig. 6D**). The group of significantly down-regulated genes included four of the 49 core-hypoxia responsive genes, likely expressed in cells physiologically exposed to low oxygen tension ^2^ (**Table SX**). The *erfVII* mutant seemed to modulate gene expression as if experiencing a mild oxidative stress under control conditions, as demonstrated by the separation of samples in the dimension primarily affected by TBHP (**Figure 6C**). Moreover, exposure to TBHP exerted a clearly distinct effect on the two genotypes. Indeed, pairwise comparisons revealed that the *erfVII* mutant induced a higher number of genes than the wild type (**Figure 6D**). We found 282 genes that were both up-regulated in the *erfVII* mutant upon TBPH treatment compared to the wild-type and down-regulated in the wild-type upon TBHP treatment compared to mock conditions. This set of genes included those involved in salicylates-mediated response, senescence, hypoxia and oxidative stress responses (**Figure 6E**). This indicates that TBHP-stabilised ERFVIIs repress these transcriptional responses. In contrast, 136 genes (18%) were not up-regulated by TBHP treatment in the *erfVII* mutant, opposite to what we observed in the wild type, indicating that TBHP-stabilised ERFVIIs also act as activators of expression of this sub-set of genes (**Figure 6D**). Additionally, the 142 genes which were down-regulated upon TBHP treatment in *erfVII* seedlings, but not in the wild-type, were enriched in GO terms related to plastids, chlorophyll biosynthesis and photosynthesis (**Figure 6F, G**). Overall, our results expand the molecular role of ERFVIIs as both activators and repressors of transcription (in a gene-selective manner) in response to hypoxia and oxidative stress, providing a potential rationale for their implication in responses to both reoxygenation and a broader range of biotic and abiotic stresses.

## Discussion

Plant flood survival requires tolerance to both submergence-induced hypoxia and the reoxygenation stress associated with desubmergence. The molecular response to submergence-induced hypoxia is well-characterised, comprising PCO/O_2_-dependent regulation of ERFVII stability and the consequent upregulation of genes enabling adaptive metabolic reconfiguration ^3,9,32,33^. What is less well characterised is the impact on this oxygen-sensing machinery of the ROS burst known to be associated with reoxygenation.^12–18^ We therefore set out to determine whether there is any cross-talk between reoxygenation, ROS and PCO-ERFVII function. We first confirmed that ERFVIIs have a role in facilitating both tolerance to, and recovery from, hypoxia-reoxygenation stress (**Figure 1**). We then confirmed that ERFVIIs persist in the nucleus after reoxygenation (**Figures 2**, **3**) and collected evidence that this is caused by H_2_O_2_-mediated inactivation of the plant Nt-Cys-degron pathway at the PCO level, leading to ERFVII stabilization (**Figure 4**). Surprisingly, we found that ROS-stabilised ERFVIIs do not promote, but rather repress, the expression of hypoxia responsive genes (**Figure 5**). Finally, we uncovered that ERFVIIs modulate transcription of a different set of genes in response to ROS than they do in response to hypoxia (**Figure 6**) suggesting the PCO-ERFVII signalling pathway can be redirected in a stress-specific manner. ERFVII transcription factors have previously been proposed to mediate responses to multiple stresses, including high salinity, heat and temperature ^28–30^. These stresses are proposed to impact the degradation of ERFVIIs via the N-degron pathway, for example salt stress is reported to decrease NO production ^30,49^ and is speculated to reduce ATE or PRT6 function ^36^. Our data, showing that exogenous supply or endogenous production of H_2_O_2_ results in PCO inhibition and ERFVII stabilisation, suggests that other conditions that involve enhanced H_2_O_2_ accumulation, such as cold stress ^4^ or pathogen attack ^50,51^, may also invoke ERFVII stabilisation via this method.

PCOs are able to respond sensitively to O_2_ availability through the kinetic effect on ERFVII oxidation. The response of PCOs to the presence of H_2_O_2_ however is less finely-tuned; inhibition of PCO function by H_2_O_2_ appeared to occur in a less specific manner, likely via a combination of both oxidation of the active site Fe(II) to Fe(III), as detected by EPR, and via oxidation of Cys residues in or near the active site, as detected by Cys-sulfinate detecting probes and tryptic digest-mass spectrometry. It is also possible that other oxidative modifications took place that we were unable to detect. These modifications are all capable of impacting substrate binding and catalysis to reduce or ablate the activity of the enzyme. While we characterised these effects *in vitro*, the sensitivity of the enzymes to H_2_O_2_-mediated inhibition (5 µM H_2_O_2_ could reduce the activity of 10 µM AtPCO4 by 50%) suggests that local concentrations of ROS at approximately equivalent concentrations to that of enzyme would be sufficient to have a biologically relevant effect. Notably, there is a precedent for ROS-mediated inhibition of oxygen-sensing enzymes in mammalian systems, were H_2_O_2_-mediated inhibition of Factor Inhibiting HIF, an Fe(II) and 2-oxoglutarate dependent oxygenase, has been reported ^52^. We were not able to observe repair of H_2_O_2_-mediated PCO inhibition with cellular antioxidants *in vitro*, however it would be of interest to examine whether modulation of redox status in cells could impact ERFVII stabilisation, or indeed whether enzymes that can reverse Cys oxidation may have a role in protecting PCOs from ROS-mediated damage.

Our intriguing findings that upon ROS treatment, despite ERFVII stabilisation and retention at HRPEs, HRGs are ‘turned off’ while responses to oxidative stress are ‘turned on’, provide an explanation for the role of ERFVIIs as a major hub to control adaptation to different adverse environment conditions ^30^. The versatility of ERFVIIs in producing stress-tailored responses is surprising and can be explained by their interaction with distinct proteins and DNA motifs. Our results indicate that oxidative stress causes only mild dislocations of ERFVII from hypoxia responsive genes and more likely turns ERFVII from positive to negative regulators of gene transcription. This could explain the limited overlap between the transcriptional changes observed between hypoxia and other stresses that involve ERFVII stabilisation, despite the crucial role these TFs play in enabling tolerance. A systematic survey of their protein and DNA partnerships under different conditions will reveal the complex regulatory networks orchestrated by these TFs. The trihelix TF HRA1, previously reported to associate with RAP2.12 and reduce its trans-activation capacity, is a promising candidate for such a role.The different assortment of conserved motifs identified in ERFVIIs in vascular plants suggests a plethora of different interactions depending on the group member. For example, the Hypoxia Responsive ERFVIIs HRE1 and HRE2 seem to play a minor role in the early activation of hypoxic genes, although it is possible they are majorly involved in other transcriptional responses ^32^. The generation of ERFVII variants devoid of specific domains will support the hunt and the characterisation of these interactions.

The ability of ERFVII to regulate different subsets of genes in response to different stimuli raises questions about the evolution of these TFs. Was the last common ancestor of the ERFVIIs, which likely appeared at the origin of land plants ^48^, a generic regulator of stress tolerance required to cope with an environment of increasing ROS? Or rather did the ERFVIIs evolve first as signaltransducersfor hypoxia, and only later expand their role to accommodate the response to other stresses? In today’s environment, however, it appears that the PCO-ERFVII sensing and signalling nexus is more nuanced than a simple response to fluctuations in O_2_ availability. Its sensitivity to oxidative stress, including that which occurs upon reoxygenation, allows dynamic control in response to a range of cues. The overall impact of this ability to switch between subsets of genes explains the key role played by ERFVIIs in plant survival of both submergence and desubmergence.

## Acknowledgements

We gratefully acknowledge the help of Elisabete Pires for assistance with conducting LC-MS/MS experiments. We also extend our gratitude to Professor Kate Carroll, Department of Chemistry and Biochemistry, Florida Atlantic University, for generously providing us with the DiaAlk probe. This work was supported by the European Research Council (ERC) under the European Union Horizon 2020 Research and Innovation Program (Grant 864888 supported S.A., D.M.G. and E.F.; Grant 101001320 supported F.L., V.S., L.D.C. and D.Z.). M.P was supported by funding from the Biotechnology and Biological Sciences Research Council (UKRI-BBSRC) [grant number BB/T008784/1]. B.F. was financially supported by the Erasmus+ programme.

## Author Contributions

S.A., M.P., E.F., B.G. and F.L. conceived and designed the study. S.A., M.P., M.L-P., B.F., L.D.C., V.S., Y.T., D.Z., W.M. and D.M.G. conducted experiments. All authors analysed data and prepared figures. S.A., M.P., E.F. and F.L. wrote the manuscript with input from all authors.

## Supplemental information

Document S1, containing Figures S1-S7 and Tables S1-S5.

Document S2, excel file containing fold change data of RNA sequencing.

## STAR Methods

### Experimental model and subject details

#### Plant materials and growth conditions

*A. thaliana* Columbia-0 (Col-0) was used as wild-type ecotype. The genotypes RAP2.12_1-28_-fLUC, RAP2.12-GFP and Δ13RAP2.12-GFP have been previously described ^3,32^. Seeds were sown in a soil:vermiculite 3:1 ratio mixture, stratified at 4 °C in the dark for 3 days, then germinated at 22 °C/20 °C with a 16h:8h, light:dark photoperiod and 100 μmol photons m^-2^ s^-1^ intensity. For *in vitro* propagation, seeds were sterilised using 70% ethanol for 1 min, incubated in 10% sodium hypochlorite (NaClO) for 10 min, followed by six washes in 1 mL sterile distilled water. Growth in liquid medium was performed inoculating 100 μL of seeds suspension, corresponding to 20–40 seeds, in 1 mL of sterile half-strength MS medium (basal salt mixture 2.15 g L^−1^, pH 5.7) supplemented with 1% sucrose in each well of 6-well plates. For growth in solid media, seeds were incubated in the dark at 4 °C for 2 days and subsequently on half-strength MS ^53^ medium, supplemented with 1% (w/v) sucrose and 0.8% (w/v) agar, and grown at 22°C,16:8 day:night photoperiod 100 μmol photons m^-2^ s^-1^ intensity.. All experiments were conducted on 7 days-old seedlings.

#### Yeast strains and culture

A haploid parental strain BY4742 (Matα; his3-Δ1; leu2-Δ0; lys2-Δ0; ura3-Δ0) was co-transformed following the LiAc/SS carrier DNA/PEG method ^54^ with PCO4-pAG415GPD and different versions of the DLOR-pAG413GPD (C-, D- or R-DLOR). All plasmids used for yeast expression were produced in previous works ^55^. Prior to transformation, cells were grown at 30°C on YPDA (20 g L-1 peptone, 10 g L-1 yeast extract, 20 g L-1 of glucose (Duchefa) and 20 mg L-1 adenine hemisulfate (Sigma-Aldrich), supplied with 20 g L-1 agar (Duchefa) when necessary). Transformants were selected on SD medium, containing 6.7 g L −1 Yeast Nitrogen Base (DIFCO), 1.37 g L−1 Yeast Dropout Medium (Sigma-Aldrich) and 20 g L−1 glucose, plus supplements (0.16 M uracil, 0.8 M histidine– HCl, 0.8 M leucine and 0.32 M tryptophan (Sigma-Aldrich) when complete), with 20 g L-1 agar when solid.

#### Bacterial strains

Bacterial strain *Escherichia coli* BL21 (DE3) was used for expression of recombinant Plant Cysteine Oxidases (PCOs). Bacteria were cultured in 2YT medium at 37°C until OD_600_ reached 0.6, protein expression was induced by addition of 0.8 mM Isopropyl-*β*-D-thiogalactoside (IPTG, Sigma-Aldrich) at 18°C for 16 h with 170 rpm in shaking incubator (Eppendorf, UK).

### Method Details

#### Construct generation

For the generation of 35S:RAP2.3-nLuc construct, a 806nt long synthetic string containing Arabidopsis codon-optimised nLuc sequence including RAP2.2 intron (**Table S1**) was synthesised in the pMK-RQ backbone by Geneart (Thermo Fisher Scientific). The destination vector pK7GWnL2 was generated as ligation between pK7GW2 ^56^ and GWnLuc-intron after restriction using XbaI and MluI (Thermo Fisher Scientific). Arabidopsis RAP2.3 CDS was amplified from Col-0 cDNA without stop codon (RAP2.3Δstop) introducing overlapping *AttB* sites using by PCR using GoTaq® DNA polymerase (Promega). Entry clone vector was then generated as a BP reaction between RAP2.3 CDS PCR product and pENTR/D-TOPO (Life Technologies). The resulting entry vector was finally recombined into the hereby generated pK7GWnL2 destination vector using the LR clonase mix II (Thermo Fisher Scientific). Primers for RAP2.3Δstop cloning and screening are listed in **Table S2**. For the generation of 35S:RAP2.3-GFP construct, entry vector containing RAP2.3 CDS was recombined with pK7GW2F ^56^ destination using the LR clonase mix II (Thermo Fisher Scientific).

For HRPE-nLuc construct design, DNA containing the SacI-attR1-ccdB-attR2-NanoLuc-HindIII sequence (**Table S3**) was *de novo* synthesized by GeneArt service (Thermo-Fisher Scientific, Waltham, MA, USA) and cloned into the pBGWL7 Gateway destination vector ^57^ The entry vector containing the HRPE:5’-UTR ADH1 sequence ^47^ was recombined into the destination vector by Gateway cloning.

For recombinant protein production, the five *pco* genes from *A. thaliana* had previously been cloned into the NdeI and XhoI sites of pET28a (Novagen), transformed into *E.coli* NEB5α competent cells (New England Biolabs) and sequences validated by Sanger sequencing (Source Biosciences) as previously described ^58^.

#### *A. thaliana* transformation

*Agrobacterium*-mediated transformation was performed to obtain RAP2.3-nLUC, RAP2.3-GFP and HRPE:nLuc stable transgenic line using the floral dip medium as previously described ^59^. T_0_ seeds were selected for resistance on agarized half-strength MS medium supplemented with the corresponding antibiotic and subsequently transferred in soil. The presence of the transgene was detected by PCR using GoTaq® DNA polymerase (Promega). T_3_ generation plants were used for the experiments.

### Chemical treatment

Oxidative stress in plants was induced by 1mM TBHP treatment diluted in MilliQ water for 6h in normoxia and 6h in hypoxia. After 3h, the media was replenished with TBHP.

### Low oxygen and reoxygenation treatments

For hypoxia treatments, seedlings were grown in six-well plates in liquid media and subjected to anaerobic conditions inside the Hypoxic Workstations (Whitley) continuously flushed with an artificial humidified atmosphere containing a mixture of oxygen (1%) and nitrogen gases (99%) at 22°C for 6 h. For sever hypoxic treatment, seedlings were grown vertically in square plates and treated with 0.5% O_2_ v/v O_2_/N_2_ for 24 h. During the hypoxic and anoxic treatments, the seedlings were maintained in the dark to avoid oxygen release by photosynthesis. Seedlings used for control samples were maintained in aerobic conditions (21% O_2_ v/v O_2_/N_2_) at dark for an equal amount of time. After low oxygen, plants were transferred to the aerobic growth conditions for reoxygenation treatment.

### Yeast treatments

For TBHP treatments, colonies were inoculated in 5mL of -his -leu SD medium, grown overnight, diluted to half in fresh media and further diluted to OD600 = 0.1. Cultures were grown for 5 h prior to the treatment. TBHP was then supplied for up to 30 min at different concentrations (0, 0.25, 0.5 and 0.75 mM). 50 µl of culture were harvested for luciferase assays, centrifuged at 15.000 rpm for 5 min, frozen and extracted in 50 mL of PLB. Luciferase was measured using the Dual-Luciferase® Reporter (DLR™) Assay System (Promega) as described before ^36^ 1 mL of culture was used for OD600 spectrometric measurements.

### Root length and survival rate measurements

For reoxygenation tolerance, 7-day old seedlings were treated as previously mentioned and primary root length and after 4 days of recovery, primary root length, fresh weight and survival rate were measured. Transparent squared plates containing Arabidopsis seedlings were scanned using the EPSON Perfection V750 PRO scanner with a 720 dots per inch resolution.

Growth rate was measured as increase in length of the primary root divided by days of recovery. Primary root length was assessed using ImageJ ^60^.

### ROS staining and quantification

Reactive oxygen species were visualised using 3,3’-diaminobenzidin (DAB, Fluorochem Ltd) staining to detect H_2_O_2_ and superoxide, respectively, using methods described previously ^61^ with minor modification. Seedlings were incubated with either 1mg/mL DAB, vacuum infiltrated for 5 min and incubated for 4-5 h at dark shaking. After staining, seedlings were washed with distilled water and bleached in several washes of 70% ethanol. 5-8 seedlings were analysed per condition using the Leica M165C stereo microscope with 2.5x magnification, followed by quantification of pixel intensity using ImageJ ^60^.

### Confocal imaging

7-day old seedlings were used for GFP detection after treatment. For nuclear localization, seedlings were stained in PBS containing with 1µg/mL 1 μg/mL 4′,6-diamidino-2-phenylindole (DAPI, Thermo Fisher Scientific) and washed three times in PBS. Imaging was performed using ZEISS LSM 880 Airyscan microscope (Department of Biology, University of Oxford), equipped with a 25x objective lens, upon laser excitation at 405 nm and collection at 410-495 nm, for DAPI imaging, and excitation at 488 nm and collection at 498-560 nm, for GFP imaging.

### Luciferase assay

Total proteins were extracted in passive lysis buffer (Promega). Firefly Luciferase activities were measured using the ONE-Glo Luciferase Assay kit (Promega) and the Nano-Glo® Luciferase Assay System (Promega) was used to measure the activity of NanoLuc® luciferase enzyme. Luciferase signal was normalized based on total protein concentration using Bradford assay ^62^.

### Recombinant protein production

Expression and His6-tag affinity purification of *At*PCOs were as previously described ^58^. Following affinity purification, the His6-tag was cleaved using TEV protease and the cleaved tag removed using a HisTrap HP column (GE Healthcare). Proteins were then purified with a HiLoad 26/600 Superdex 75 prep grade size exclusion column (GE Healthcare) equilibrated with 50 mM Tris (pH 7.5) and 0.4 M NaCl. Protein purity was assessed with SDS-PAGE.

### *In vitro* H_2_O_2_ oxidation assay of PCOs

10 μM of recombinant PCOs enzymes (PCO 1 to 5) were incubated with H_2_O_2_ or equal volume of H_2_O in 50 mM HEPES buffer at pH 7.4 (herein termed reaction buffer) at 4°C for 30 minutes. Excess H_2_O_2_ was removed using a Micro Bio-Spin P-6 chromatography column (Bio-Rad), equilibrated with reaction buffer. 200 μM RAP2_2-15_ peptide (CGGAIISDFIPPPR, purchased from GL Biochem (China)) was reacted with 1 μM PCO (H_2_O_2_ treated or non-treated) at 25 °C for the required time before 5 μL samples were quenched in 45 μL of 5% formic acid to stop the enzymatic reaction. Peptide masses were subsequently analysed using an Agilent RapidFire RF360 sampling robot connected to an Agilent 6530 Accurate-Mass Q-ToF mass spectrometer operated in positive electrospray mode. Product distributions were assigned via the relative integrated areas of the peaks corresponding to products of interest. Spectra were visualised on Qualitative Analysis (Version B.07.00) and Agilent RapidFire Integrator (Version 4.3.0.17235) was used to calculate integrated peak areas.

### H_2_O_2_ oxidation assay of RAP2_2-15_ peptide

200 µM RAP2_2-15_ stock solutions were prepared in reaction buffe and treated with 400 µM freshly prepared dithiothreitol (DTT) for 45 minutes at room temperature (25 °C) to ensure peptides were monomeric (in all experiments unless otherwise mentioned). Peptides were then treated with H_2_O_2_ under conditions required by the experiment. Time-course experiments were conducted in 2 mL deep welled plates and analysed in real time using RapidFire mass spectrometry as described above.

### Peptide fragmentation by LC-MS/MS

20 µM of DTT reduced RAP2_2-15_ was treated with 1mM H_2_O_2_ at room temperature for 1h. 5 µL sample was diluted with 45 µL 5% formic acid for measurement. LC-MS/MS was carried out using an Acquity UPLC system coupled to a Xevo G2-XS Q-ToF mass spectrometer on a Chromolith Performance RP-18e 100-2 mm HPLC column (Merck) at 40 °C as above. Ions with a m/z ratio of 1474.7 (+32 Da of AtRAP2_2-15_) were selected for sequential fragmentation under a collision energy of 80V. Fragment assignment was made via comparison of the obtained spectrum with computationally predicted fragment patterns, calculated using the University of California, San Francisco webpage tool Protein Prospector (Version 6.3.1).

### Protein analysis

Protein mass measurement: 100 μM of recombinant AtPCO4 enzyme was incubated with 1 mM H_2_O_2_ or equal volume of H_2_O in reaction buffer for 1h at 25°C. Excess H_2_O_2_ was removed using a Micro Bio-Spin P-6 chromatography column (Bio-Rad) equilibrated with reaction buffer. Total mass of the H_2_O_2_ treated or nontreated AtPCO4 was measured using RapidFire mass spectrometry as described above.

Cysteine oxidation detection: After passing the column, H_2_O_2_ treated or nontreated AtPCO4 were incubated with 1 mM BioDiaAlk in dark for 1 h at 25°C, followed by 10 mM DTT reduction for 1h at 25°C. Equal amount proteins were separated on SDS-PAGE, transferred into polyvinylidene difluoride (PVDF) membrane followed by streptavidin-HRP blot at 1:1000 dilution or anti-his-HRP blot at 1:10,000 dilution, the protein signal was visualized by chemiluminescence (ECL Plus, Pierce).

### LC-MS/MS data acquisition

15 μg recombinant AtPCO4 enzyme was incubated with H_2_O_2_ or equal volume of H_2_O in reaction buffer for 1h at 25°C. After removing excess H_2_O_2,_ the enzyme was reduced with 85 mM DTT in 50 mM ammonium bicarbonate (Ambic) for 40 min at 56° C, followed by 55mM Iodoacetamide (IAA) incubation in 50 mM Ambic for 30 min in the dark at RT. To eliminate excess IAA, samples were reduced again with 85 mM DTT in 50 mM Ambic for 10 min in the dark at RT. In-solution trypsin digestion was performed by adding trypsin in a 1:50 (W/W) ratio overnight at 37 °C, followed by desalting using C18 ZipTip. The resulting tryptic peptides were resuspended in 40 μL of Milli-Q water with 2% acetonitrile and 0.1% formic acid, and 2ul were analysed on a NanoAcquity-UPLC system (Waters) connected to an Orbitrap Elite mass spectrometer (Thermo Fischer Scientific) possessing an EASY-Spray nano-electrospray ion source (Thermo Fischer Scientific). The peptides were trapped on an in-house packed guard column (75 μm i.d. x 20 mm, Acclaim PepMap C18, 3μm, 100 Å) using solvent A (0.1% Formic Acid in water) at a pressure of 140 bar. The peptides were separated on an EASY-spray Acclaim PepMap® analytical column (75 μm i.d. × 50 mm, RSLC C18, 3 μm, 100 Å) using a linear gradient (length: 100 minutes, 3 % to 60 % solvent B (0.1% formic acid in acetonitrile), flow rate: 300 nL/min). The separated peptides were electrospray directly into the mass spectrometer operating in a data-dependent mode using a CID-based method. Full scan MS spectra (scan range 350-1500 m/z, resolution 120000, AGC target 1e6, maximum injection time 250 ms) and subsequent CID MS/MS spectra (AGC target 5e4, maximum injection time 100 ms) of 10 most intense peaks were acquired in the Ion Trap. CID fragmentation was performed at 35 % of normalised collision energy, and the signal intensity threshold was kept at 500 counts. The CID method used performs beam-type CID fragmentation of the peptides.

### Processing data

The analysis was performed with Peaks 8.5. The raw MS file was searched against the TAIR database. Trypsin with a maximum of 3 missed cleavages and one unspecific end was selected as the protease. Carbamidomethylation (Cysteine) was set as a fixed modification, and oxidation (Methionine) and deamination (Asparagine, Glutamine) were set as variable modifications. Precursor mass tolerance was set as 15 ppm. Fragment mass tolerances for CID were set to 0.8 Da, respectively. All spectra were manually validated. All peptides present at −10lgP> 20, and spectra were manually checked, validated, or disqualified.

### Electron paramagnetic resonance (EPR) of AtPCO4

EPR spectroscopy was performed on a Bruker EMXmicro spectrometer with a Premium bridge connected to an ER4122SHQE-W1 cavity fitted to an Oxford Instruments ESR900 cryostat. The microwave source operated at 9.3891(17) GHz. The spectra were recorded at 10 K with liquid helium cryogen. Protein (200 µM) and control solutions were frozen in liquid N_2_. Spectra were obtained as two 5 min scans from 10 to 650 mT using time constant 20.48 ms, a microwave power of 200 uW modulation amplitude of 1 mT, and a modulation frequency of 100 kHz.

### RNA extraction and real-time qPCR analyses

Total RNA was extracted from 60-80 mg of plant material using the phenol-chloroform extraction method as described previously ^3^ RNA concentration was quantified using a NanoDrop ND-1000 (Thermo Scientific) and RNA integrity was tested on 1% agarose gel. One microgram of total RNA was subjected to DNase Treatment (Thermo Scientific™) and retro-transcribed using qPCRBIO cDNA Synthesis Kit (PCR Biosystems). Real-time quantitative PCR was performed with StepOnePlus™ Real-Time PCR System using a *Power* SYBR™ Master Mix (Thermo Fisher Scientific™). *Ubiquitn10* (*AT4G0532*) was used as housekeeping genes for Arabidopsis analysis. Four biological replicates were extracted for each condition, each represented by two technical replicates and the average expression was calculated. The primer pairs used in the real-time qRT-PCR are listed in **Table S4**. Relative expression of each individual gene was calculated using the 2−Ct method ^63^.

### Chromatin Immunoprecipitation Assay

ChIP was performed using a modified version of ^64^ Chromatin was extracted from 2 g of 7-day-old Δ13RAP2.12-GFP seedlings grown in sterile liquid half-strength MS medium, supplemented with 1% w/v sucrose, under controlled conditions (16h:8h light:dark photoperiod, at 22 °C). Seedlings were treated with 1 mM TBHP, or DMSO as control, in 1 mL of fresh liquid 1% w/v sucrose half-strength MS medium for six hours in the dark. After chromatin sonication, a volume corresponding to 2% of the total lysate volume was set aside to be used as ‘input’ sample. 5 µg of GFP antibody (Roche, cat. No. 11814460001) were used for immunoprecipitation and the antibody was pulled down from the nuclear lysate after sonication using Dynabeads™ Protein G magnetic beads (Thermo Scientific). At the end of the reverse cross-link step, DNA was purified using QIAquick PCR Purification Kit (Qiagen) and eluted in a final volume of 30 µL. Enrichment of genes in the chromatin immunoprecipitate was detected through real-time qPCR (CFX384 Touch Real-Time PCR Detection System, BioRad) with a triple technical replicate for each of the four biological replicates, applying the percent input method. To calculate the ratio between immunoprecipitated (ip) DNA and input DNA, Log250 was subtracted from the raw Ct values of the input, before the Ctip-input value was obtained. The final enrichment was calculated as 2^-ddCt. The primer sequences used are listed in **Table S5**.

### RNA sequencing

Col-0 and *erfVII* seedlings were grown for 7 days in a six-well plate in liquid media, followed by 6 h treatment with 1 mM TBPH, at dark, or mock. At the end of the treatment, samples were harvested and frozen in liquid nitrogen. RNA was isolated using GeneJET RNA Purification Kit (Thermo Scientific™) as per manufacturer’s instruction. RNA sequencing was exploited in paired-end mode using the Illumina Sequencing PE150 on the NovaSeq 6000 platform (Novogene). Transcriptomic analyses were conducted in R software (version 4.3.1). After quality check using FastQC, reads were aligned on *Arabidopsis thaliana* full genome (TAIR 10) using *Rsubread* ^65^ version 2.16.1) and counted using *featureCounts* software ^66^ Multi-Dimensional scaling (MDS) plot was used to assess similarities and differences between samples based on their gene expression profiles). Differentially expressed genes were identified using *edgeR* ^67^ version 3.42.4). Differentially expressed genes with ≥ |1.5| expression fold change and FDR <0.05 (**Table S6**) were selected for subsequent analysis. GO term enrichment analysis of the differentially expressed genes was conducted using *clusterProfiler* ^68^(version 4.10.1).

### Statistical analyses

Statistical analyses were performed using GraphPad Prism 10.2.3(403) and R Statistical Software (version 4.3.1, Foundation for Statistical Computing, Vienna, Austria). Normal distribution and homogeneity of variances of data were evaluated through Shapiro test and Levene’s test, respectively. Student’s t-test, ANOVA or Kruskal-Wallis test followed by Tukey HSD post hoc test (p < 0.05) were performed to establish the statistical significance of differences. Additional information is provided in figure legends.

### Additional Software

Molecular structures and reaction schemes were drawn using ChemDraw Professional (Version 21.0.0.28). Graphs were made and annotated using GraphPad Prism 10.2.3(403) and R Statistical Software (version 4.3.1, Foundation for Statistical Computing, Vienna, Austria).

## Data availability

RNA sequencing raw data generated for this study has been deposited in the Sequence Read Archive (SRA) at the National Centre for Biotechnology Information under BioProject ID PRJNA1171625. Any additional information required to reanalyze the data reported in this paper is available from the lead contacts upon request.

## Materials availability

All unique/stable reagents generated in this study are available from the lead contacts without restriction.

