## Supplementary material for "H_2_O_2_ repurposes the plant oxygen-sensing machinery to control the transcriptional response to oxidative stress": S1: Supplementary data


### Authors/affiliations

Salma Akter[1†], Monica Perri[2†], Mikel Lavilla-Puerta[2], Beatrice Ferretti[2,3], Laura Dalle Carbonare[2], Vinay Shukla[2], Yuri Telara[2], Daai Zhang[2], Dona M. Gunawardana[1], William K. Myers[1], Beatrice Giuntoli[4], Emily Flashman[2*] and Francesco Licausi[2**]

### Author list footnotes

[1] Department of Chemistry, University of Oxford, Mansfield Road OX1 3TA, Oxford, UK

[2] Department of Biology, University of Oxford, South Parks Road OX13RB, Oxford, UK

[3] University of Bologna, Via Zamboni, 33 40126, Bologna, IT

[4] Department of Biology, University of Pisa, Via Luca Ghini 13, Pisa, IT

[†] Equal contribution,

[*] Corresponding authors

### Contact info

**Figure S1. RAP2.3 localizes to the nucleus upon oxidative stress and reoxygenation in a time-dependent manner**

(**A**) Stabilization of 35S:RAP2.3-GFP (green) in 7-day old Arabidopsis seedlings upon 1 mM TBHP or mock treatment in normoxia (21% $O_2$), hypoxia (1% $O_2$) or after 3 h of reoxygenation (scale bar, 50 µM). (**B**) Time-dependent stabilization of 35S:RAP2.3-GFP (green) in 7-day old seedlings over 2 h of 1 mM TBHP treatment in air (scale bar, 50 µM).

A

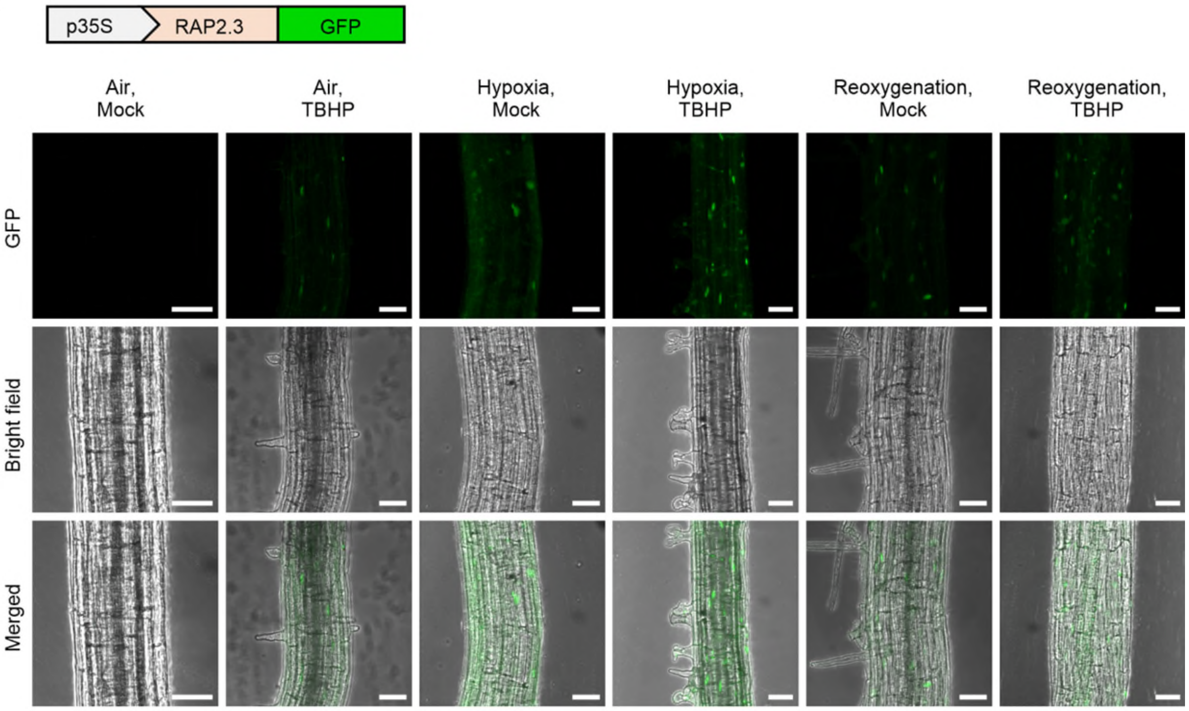


B

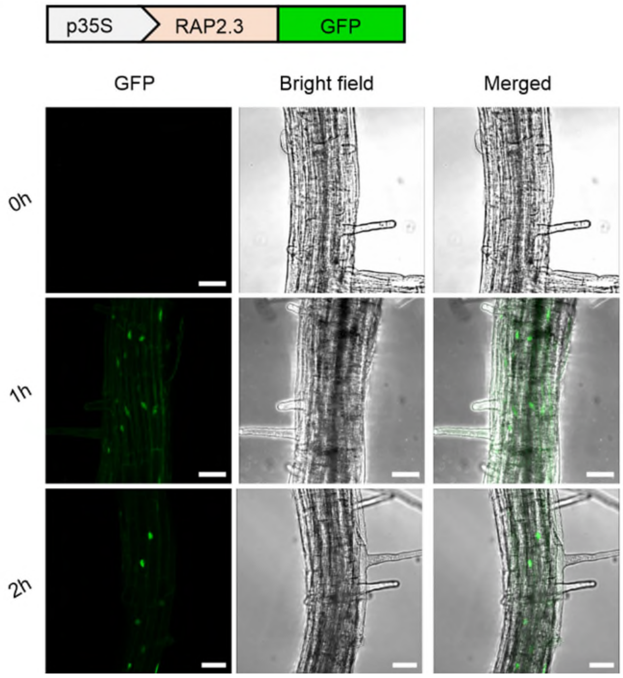

**Figure S2. Measurement of ERFVII stability upon oxidative stress in plant and yeast-based reporter systems.**

(**A**) Relative FLuc activity of 35S:RAP2.12$_{2\text{-}28}$-FLuc seedlings in air, hypoxia and reoxygenation upon 1 mM TBHP or mock treatment ($n$ = 4). (**B**) Relative FLuc activity of yeast expressing C-DLOR and PCO4 over time. (**C**) Effect of TBHP doses on C-DLOR activity ($n$ = 5). (**D**) Measurement of optical density at 600 nm (OD600) of yeast culture subjected to different doses of TBHP ($n$ = 5). Two-way (**A, C**) or one-way (**B, D**) ANOVA followed by Tukey HSD test, $p<0.05$ where different letters indicate statistically different groups.

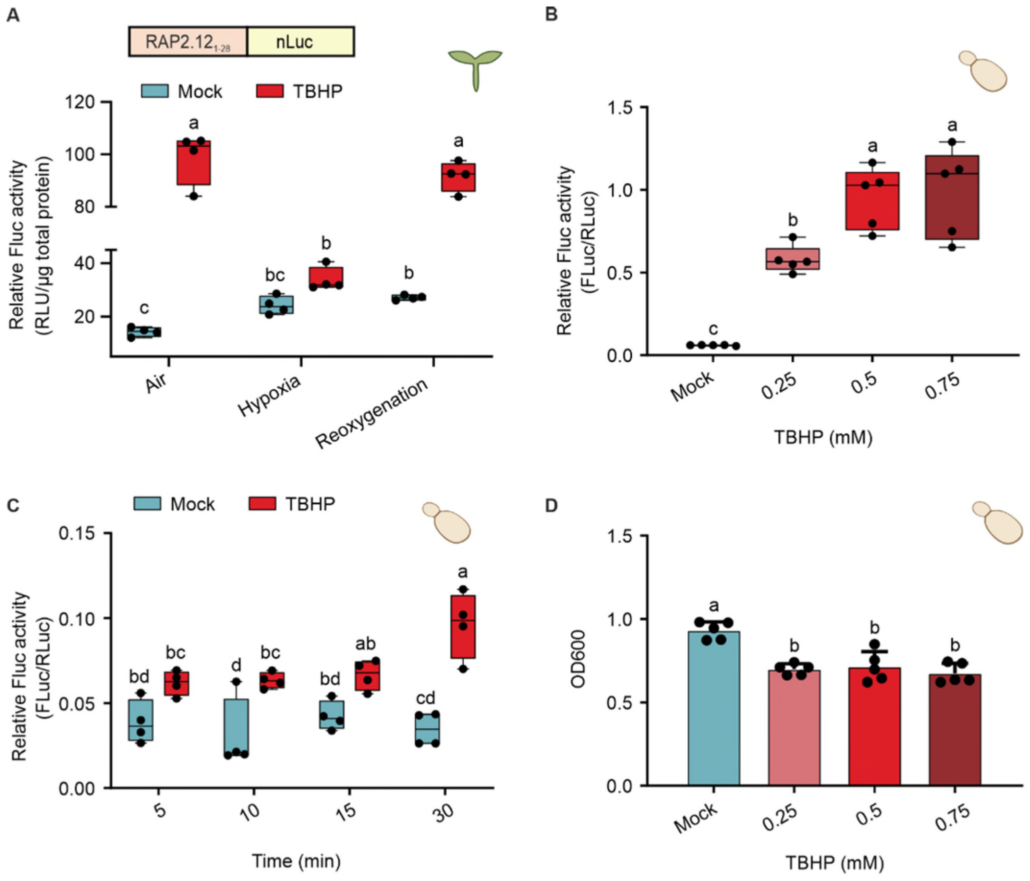

**Figure S3. Direct $H_2O_2$ oxidation of $RAP2_{2\text{-}15}$ peptide N-terminal cysteine**

(**A**) UPLC-Mass spectrometry spectra showing mass increases of $RAP2_{2\text{-}15}$ after 1 mM $H_2O_2$ treatment for 1 h (White et al., 2018) . (**B**) $H_2O_2$ causes sulfenic (SOH), sulfinic ($SO_2H$) and sulfonic acid ($SO_3H$) modifications on the N-terminal cysteine residue of $RAP2_{2\text{-}15}$ after 1 mM $H_2O_2$ treatment for 1 h. (**C**) Tandem mass spectrometry confirms $H_2O_2$-mediated oxidative modification on Nt-Cys of $RAP2_{2\text{-}15}$; Spectrum shows b ions and y ions of fragmented 1474.7 Da peptide (Nt-Cys-$SO_2H$) following $H_2O_2$ treatment $RAP2_{2\text{-}15}$ (collision energy 80 V). Expected y ions are observed, however b ions were predominantly observed with a consistent mass loss of 81 Da, termed b* ions. These ions likely correspond to loss of $SO_2$ and $NH_3$ upon fragmentation, as it has been observed previously for an oxidative modification on N-terminal cysteine (Masson et al. 2019). (**D**) Table showing the matched mass of the predicted fragments (b, b* and y) with the observed mass of $RAP2_{2\text{-}15}$ (1474.7 Da) under $H_2O_2$ treatment.

A

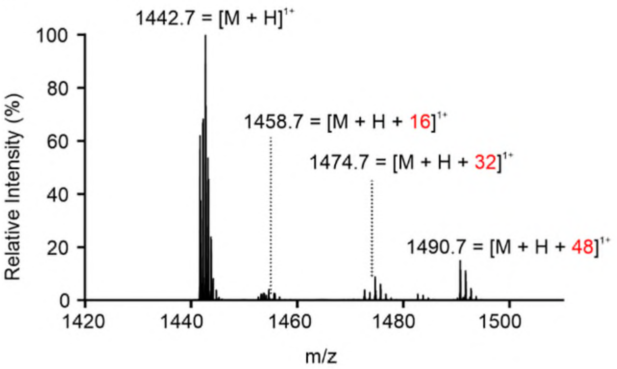


B

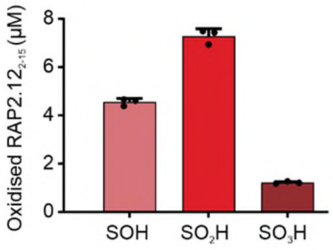


C

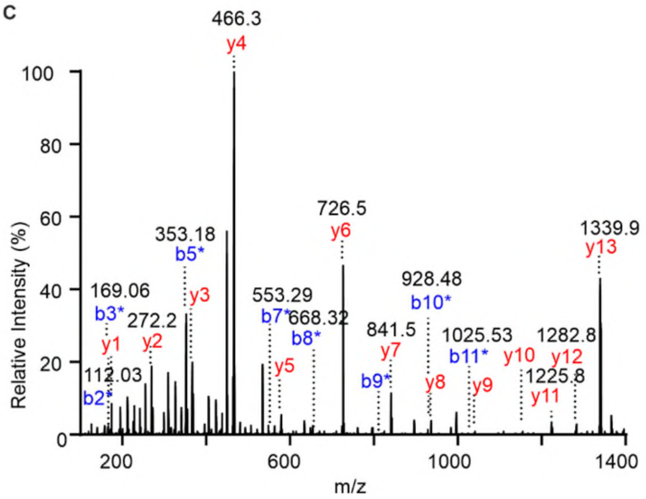


D

| b*(b- 81 Da) | b | # | Seq of $RAP2_{2\text{-}15}$ | # | y |
|---|---|---|---|---|---|
| | --- | 1 | C(+32 Da) | 14 | --- |
| 112.0 | 193.0 | 2 | G | 13 | 1339.74 |
| 169.1 | 250.1 | 3 | G | 12 | 1282.72 |
| 240.1 | 321.1 | 4 | A | 11 | 1225.69 |
| 353.2 | 434.2 | 5 | I | 10 | 1154.66 |
| 466.3 | 547.3 | 6 | I | 9 | 1041.57 |
| 553.3 | 634.3 | 7 | S | 8 | 928.489 |
| 668.3 | 749.3 | 8 | D | 7 | 841.457 |
| 815.4 | 896.4 | 9 | F | 6 | 726.43 |
| 928.5 | 1009.5 | 10 | I | 5 | 579.361 |
| 1025.5 | 1106.5 | 11 | P | 4 | 466.277 |
| 1122.6 | 1203.6 | 12 | P | 3 | 369.225 |
| 1219.6 | 1300.6 | 13 | P | 2 | 272.172 |
| | --- | 14 | R | 1 | 175.119 |

**Figure S4. $H_2O_2$ inhibits recombinant PCO enzymes.**

(**A**) Intact protein mass spectra of 100 µM $H_2O_2$ treated and non-treated 10 µM AtPCO4 measured by RapidFire mass spectrometry**.** (**B**) Streptavidin blot of BioDiaAlk labeling of sulfinic acids in PCO4. (**C**) Table showing percentage of ion (b and y) intensity of $H_2O_2$ treated and non-treated PCO4. (**D**) IAM (+ 57.02) modifications on b ion and y ion fragments of the peptide containing Cys165 of PCO4. (**E**) Sulfinic acid (+ 31.98) modifications on b ion and y ion fragments of the peptide containing Cys165 of PCO4. (**F**) Sulfonic acid (+ 47.97) modifications on b ion and y ion fragments of the peptide containing Cys165 of PCO4.

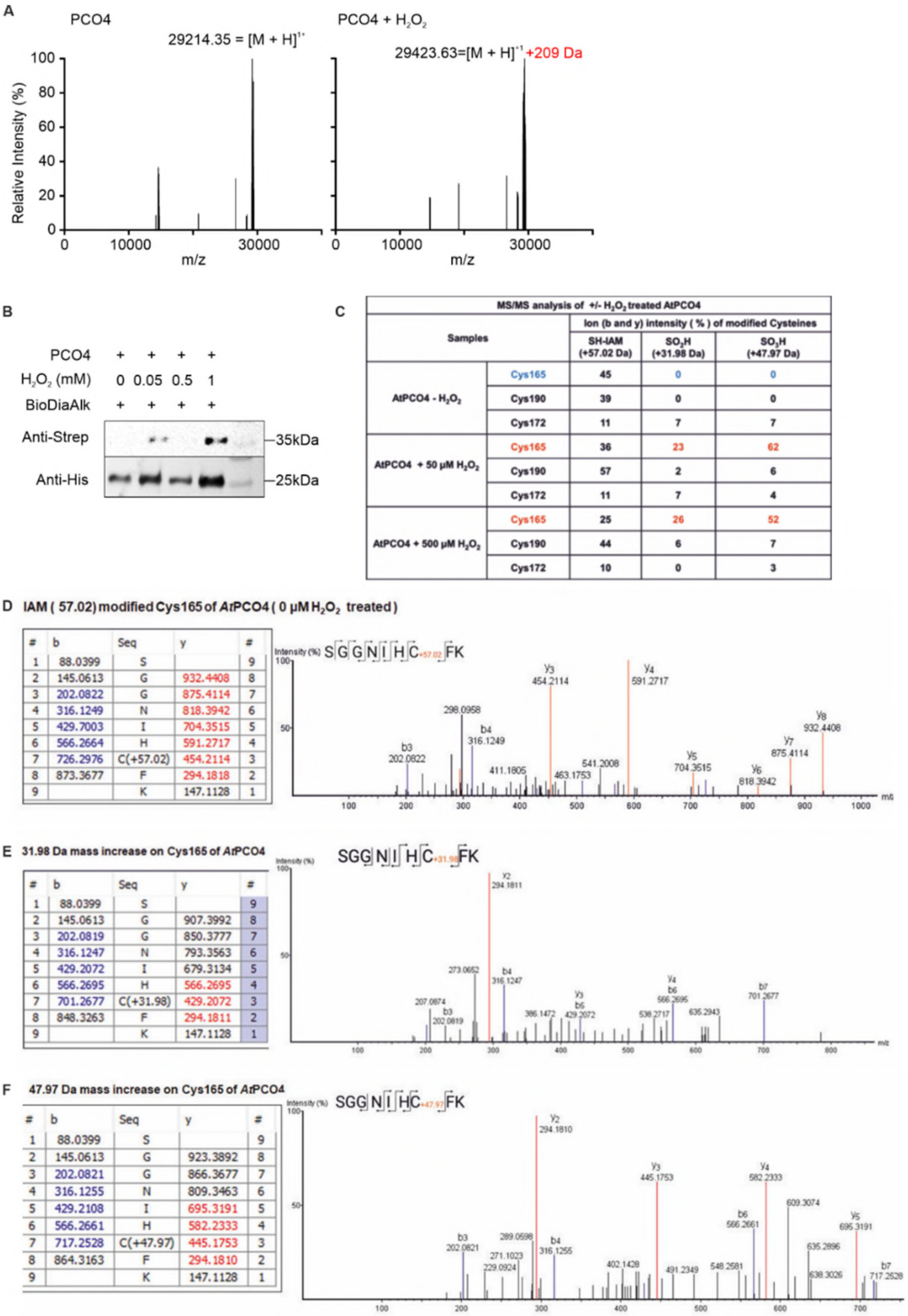


| MS/MS analysis of +/- $H_2O_2$ treated AtPCO4 | | | | |
|---|---|---|---|---|
| Samples | | Ion (b and y) intensity ( % ) of modified Cysteines | | |
| | | SH-IAM (+57.02 Da) | $SO_2H$ (+31.98 Da) | $SO_3H$ (+47.97 Da) |
| AtPCO4 - $H_2O_2$ | Cys165 | 45 | 0 | 0 |
| | Cys190 | 39 | 0 | 0 |
| | Cys172 | 11 | 7 | 7 |
| AtPCO4 + 50 µM $H_2O_2$ | Cys165 | 36 | 23 | 62 |
| | Cys190 | 57 | 2 | 6 |
| | Cys172 | 11 | 7 | 4 |
| AtPCO4 + 500 µM $H_2O_2$ | Cys165 | 25 | 26 | 52 |
| | Cys190 | 44 | 6 | 7 |
| | Cys172 | 10 | 0 | 3 |

D

| # | b | Seq | y | # |
|---|---|---|---|---|
| 1 | 88.0399 | S | | 9 |
| 2 | 145.0613 | G | 932.4408 | 8 |
| 3 | 202.0822 | G | 875.4114 | 7 |
| 4 | 316.1249 | N | 818.3942 | 6 |
| 5 | 429.7003 | I | 704.3515 | 5 |
| 6 | 566.2664 | H | 591.2717 | 4 |
| 7 | 726.2976 | C(+57.02) | 454.2114 | 3 |
| 8 | 873.3677 | F | 294.1818 | 2 |
| 9 | | K | 147.1128 | 1 |

E

| # | b | Seq | y | # |
|---|---|---|---|---|
| 1 | 88.0399 | S | | 9 |
| 2 | 145.0613 | G | 907.3992 | 8 |
| 3 | 202.0819 | G | 850.3777 | 7 |
| 4 | 316.1247 | N | 793.3563 | 6 |
| 5 | 429.2072 | I | 679.3134 | 5 |
| 6 | 566.2695 | H | 566.2695 | 4 |
| 7 | 701.2677 | C(+31.98) | 429.2072 | 3 |
| 8 | 848.3263 | F | 294.1811 | 2 |
| 9 | | K | 147.1128 | 1 |

F

| # | b | Seq | y | # |
|---|---|---|---|---|
| 1 | 88.0399 | S | | 9 |
| 2 | 145.0613 | G | 923.3892 | 8 |
| 3 | 202.0821 | G | 866.3677 | 7 |
| 4 | 316.1255 | N | 809.3463 | 6 |
| 5 | 429.2108 | I | 695.3191 | 5 |
| 6 | 566.2661 | H | 582.2333 | 4 |
| 7 | 717.2528 | C(+47.97) | 445.1753 | 3 |
| 8 | 864.3163 | F | 294.1810 | 2 |
| 9 | | K | 147.1128 | 1 |

**Figure S5. Hypoxic response in 35S:RAP2.3-nLuc seedlings.**

Relative expression of six hypoxic responsive genes (*LBD41*, *ADH1*, *PDC1*, *SUS1*, *HRA*1 and *HB1*) in 35S:RAP2.3-nLuc 7-day old seedlings subjected to air, hypoxia or reoxygenation upon TBHP or mock treatment (*n* = 4). Statistical analyses were conducted using Two-way ANOVA followed by Tukey HSD test, p<0.05. Different letters indicate statistically different groups.

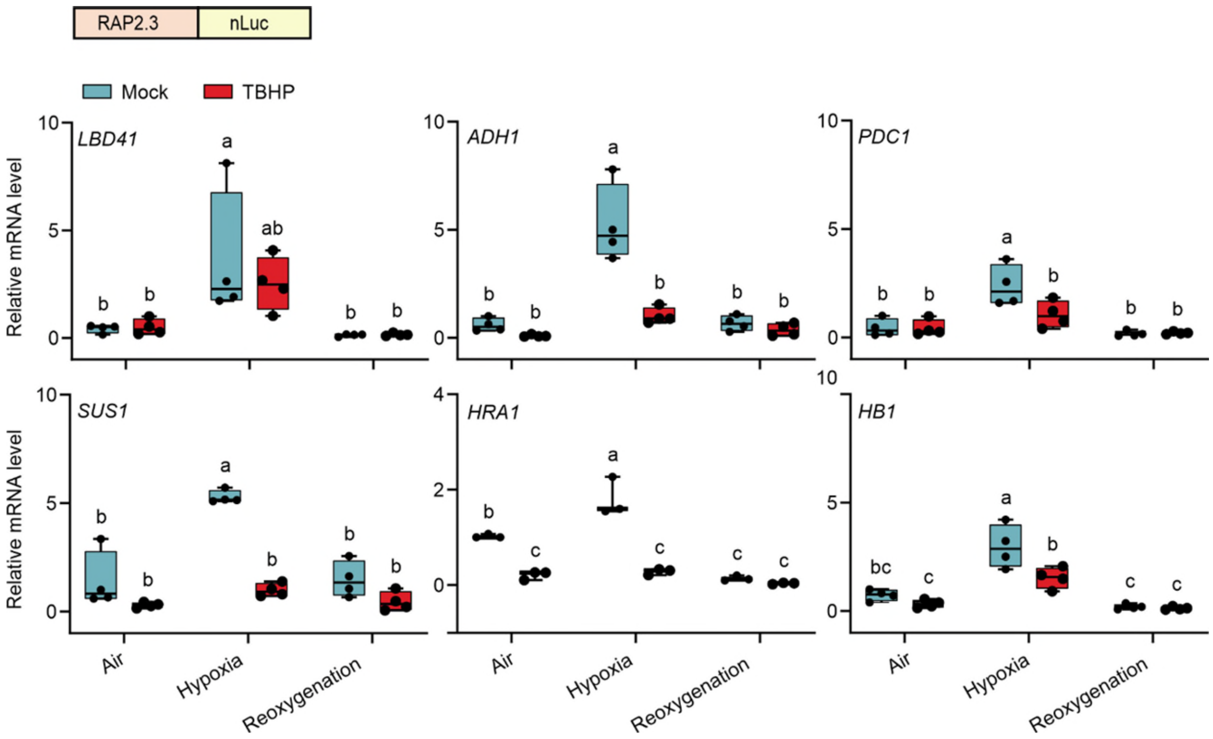

**Figure S6. Hypoxic response in HRPE:nLuc seedlings.**

Relative expression of six hypoxic responsive genes (*LBD41*, *ADH1*, *PDC1*, *SUS1*, *HRA*1 and *HB1*) in HRPE:nLuc 7-day old seedlings subjected to air, hypoxia or reoxygenation upon TBHP or mock treatment (*n* = 4). Statistical analyses were conducted using Two-way ANOVA followed by Tukey HSD test, p<0.05. Different letters indicate statistically different groups.

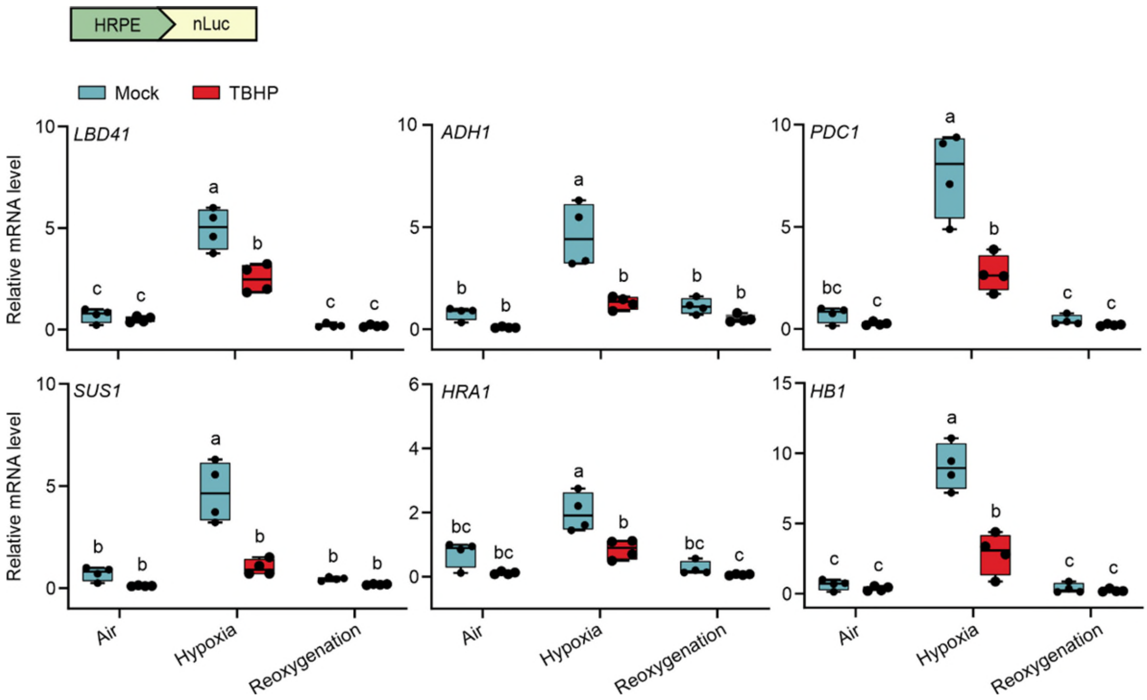

**Figure S7. Gene ontology (GO) terms analysis for biological processes of up- and down-regulated genes.**

Pairwise analysis of GO terms associated with transcripts differentially regulated in *erfVII* and wild-type seedlings, with or without TBHP treatment. Circle size indicates the gene count per GO term, with color maps indicating the False Discovery Rate (FDR) value (p.adjust).

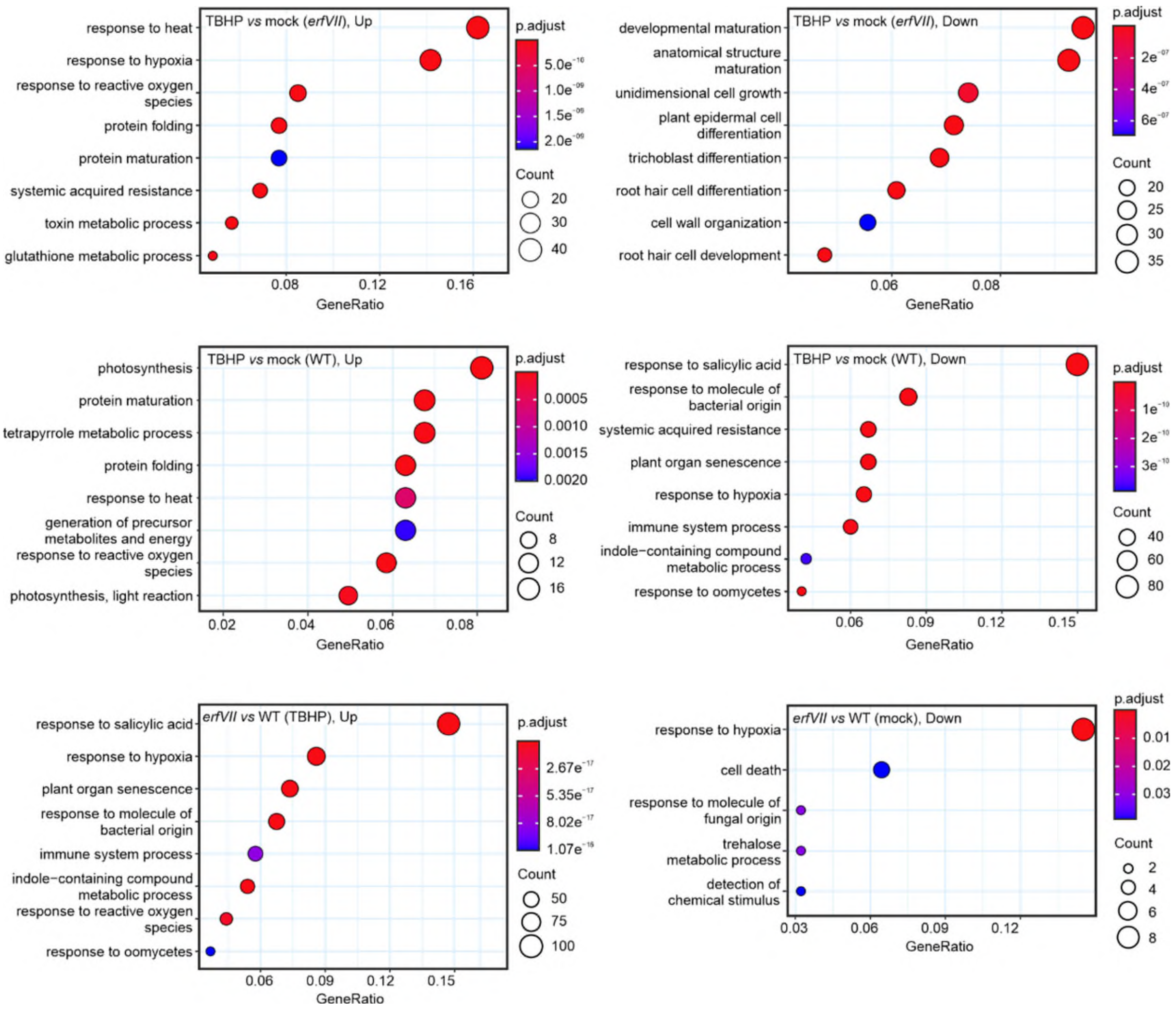

### Table S1. Sequence of Arabidopsis codon-optimised nLuc-intron synthetic sequence

ATGGTTTTCACCCTTGAGGACTTCGTTGGAGATTGGAGACAGACCGCTGGATACAACCTTGATCAGGTGTTG
GAGCAAGGTGGGGTGTCATCTTTGTTCCAGAACCTCGGAGTTAGCGTGACCCCTATCCAGAGAATCGTTCT
CTCTGGTGAGAACGGGCTCAAGATCGATATCCACGTGATCATCCCTTACGAGGGACTTAGCGGAGATCAGA
TGGGACAGATCGAGAAGATTTTCAAGGTGGTGTACCCTGTGGACGACCACCACTTCAAGGTTATCCTCCATT
ACGGtaaatatttagggattgactttagtttgagcgattgaattggttagaaattaagtttcgattctatttagctctgttcatagatacctttatctgcttacgattttgatt
gttgtataaagcatgaggtttgatttgtgtgatatttatatgcatatgaggttgagtgatgagacttgatttgtgtgatgagatttgattagagattgttgatgagttatttat
agaagaactcttttgttttgttgttgtttcactactagGAACCCTCGTGATCGATGGTGTGACCCCAAACATGATCGACTACTTCGG
TAGACCGTACGAGGGAATCGCTGTGTTCGATGGAAAGAAGATTACCGTCACTGGGACCCTCTGGAACGGGA
ACAAGATTATCGATGAGAGGCTCATCAACCCGGACGGCTCACTTTTGTTCAGAGTGACTATCAACGGTGTGA
CCGGTTGGAGACTTTGCGAGAGAATTTTGGCTGGAAGCAGCGGAGCTATCGCCATGGAATGA

### Table S2. List of primers for generation and screening of RAP2.3-nLuc and RAP2.3-GFP transgenic line

| Primer Name | Primer |
|---|---|
| RAP2.3Δstop Fw | AAAAAAGCAGGCTCCATGTGTGGCGGTGCTATTAT |
| RAP2.3Δstop Rv | CAAGAAAGCTGGGTGCTCATACGACGCAATGACAT |
| AttB1 | GGGGACAAGTTTGTACAAAAAAGCAGGCTCC |
| AttB2 | CACCCAGCTTTCTTGTACAAAGTGGTCCCC |
| nLuc Rv | AGTCCCTCGTAAGGGATGAT |
| GFP Rv | ACAACTCCAGTGAAAAGTTC |

### Table S3. Sequence of Arabidopsis codon-optimised HRPE-nLuc synthetic construct.

ctcccatatggtcgacctgcaggcggccgcactagtgatatcacaagtttgtacaaaaaagctgaacgagaaacgtaaaatgatataaatatcaatatattaaa
ttagattttgcataaaaaacagactacataatactgtaaaacacaacatatccagtcactatggcggccgcattaggcaccccaggctttacactttatgcttccg
gctcgtataatgtgtggattttgagttaggatccggcgagattttcaggagctaaggaagctaaaatggagaaaaaaatcactggatataccaccgttgatatatc
ccaatggcatcgtaaagaacattttgaggcatttcagtcagttgctcaatgtacctataaccagaccgttcagctggatattacggcctttttaaagaccgtaaaga
aaaataagcacaagttttatccggcctttattcacattcttgcccgcctgatgaatgctcatccggaattccgtatggcaatgaaagacggtgagctggtgatatgg
gatagtgttcacccttgttacaccgttttccatgagcaaactgaaacgttttcatcgctctggagtgaataccacgacgatttccggcagtttctacacatatattcgca
agatgtggcgtgttacggtgaaaacctggcctatttccctaaagggtttattgagaatatgtttttcgtctcagccaatccctgggtgagtttcaccagttttgatttaaa
cgtggccaatatggacaacttcttcgcccccgttttcaccatgggcaaatattatacgcaaggcgacaaggtgctgatgccgctggcgattcaggttcatcatgcc
gtctgtgatggcttccatgtcggcagaatgcttaatgaattacaacagtactgcgatgagtggcagggcggggcgtaaacgcgtggatccggcttactaaaagc
cagataacagtatgcgtatttgcgcgctgatttttgcggtataagaatatatactgatatgtatacccgaagtatgtcaaaaagaggtgtgctatgaagcagcgtatt
acagtgacagttgacagcgacagctatcagttgctcaaggcatatatgatgtcaatatctccggtctggtaagcacaaccatgcagaatgaagcccgtcgtctg
cgtgccgaacgctggaaagcggaaaatcaggaagggatggctgaggtcgcccggtttattgaaatgaacggctcttttgctgacgagaacagggactggtga
aatgcagtttaaggtttacacctataaaagagagagccgttatcgtctgtttgtggatgtacagagtgatattattgacacgcccgggcgacggatggtgatcccc
ctggccagtgcacgtctgctgtcagataaagtctcccgtgaactttacccggtggtgcatatcggggatgaaagctggcgcatgatgaccaccgatatggccag
tgtgccggtctccgttatcggggaagaagtggctgatctcagccaccgcgaaaatgacatcaaaaacgccattaacctgatgttctggggaatataaatgtcag
gctcccttatacacagccagtctgcaggtcgaccatagtgactggatatgttgtgttttacagtattatgtagtctgtttttttatgcaaaatctaatttaatatattgatattta
tatcattttacgtttctcgttcagctttcttgtacaaagtggtgataaaaaaaatggttttcacccttgaggacttcgttggagattggagacagaccgctggatacaacc
ttgatcaggtgttggagcaaggtggggtgtcatctttgttccagaacctcggagttagcgtgacccctatccagagaatcgttctctctggtgagaacgggctcaa
gatcgatatccacgtgatcatcccttacgagggacttagcggagatcagatgggacagatcgagaagattttcaaggtggtgtaccctgtggacgaccaccact
tcaaggttatcctccattacggaaccctcgtgatcgatggtgtgaccccaaacatgatcgactacttcggtagaccgtacgagggaatcgctgtgttcgatggaa
agaagattaccgtcactgggaccctctggaacgggaacaagattatcgatgagaggctcatcaacccggacggctcacttttgttcagagtgactatcaacggt
gtgaccggttggagactttgcgagagaattttggctggaagcagcggagctatcgccatggaatgaccgcggccatgctagagtccgcaaaaatcaccagtct
ctctctacaaatctatctctctctattttctccagaataatgtgtgagtagttcccagataagggaattagggttcttatagggtttcgctcatgtgttgagcatataaga
aacccttagtatgtatttgtatttgtaaaatacttctatcaataaaatttctaattcctaaaaccaaaatccagtgacctgcaggcatgcgacgtcgggccca

**Table S4. List of primers for qRT-PCR**

| Gene | AGI Code | Forward Primer | Reverse Primer |
|---|---|---|---|
| *UBQ10* | *AT4G05320* | GGCCTTGTATAATCCCTGATGAATAAG | AAAGAGATAACAGGAACGGAAA CATAGT |
| *LBD41* | *AT3G02550* | TGAAGCGCAAGCTAACGCA | ATCCCAGGACGAAGGTGATTG |
| *ADH1* | *AT1G77120* | TATTCGATGCAAAGCTGCTGTG | CGAACTTCGTGTTTCTGCGGT |
| *PDC1* | *AT4G33070* | CACAGAATCTTCAATGTTCTTACC | CCATGATAAAGCGTACATGGAA |
| *SUS1* | *AT5G20830* | ACGCTGAACGTATGATAACGCG | AACCCTGGAAAGCAAGGCAAG |
| *HRA1* | *AT3G10040* | ACAACCACCGCAACAGAATCC | TCTCCGCAATTCTCGCCAT |
| *HB1* | *AT2G16060* | AATGATTTATAACTGCAGGTGGC | TCATAAGCCTGACCCCAAGC |
| *CRK36* | *AT4G04490* | CCGGATGCGGAGGAGGATTT | TACCGCTATCTCTTGCCCGC |
| *GSTU24* | *AT1G17170* | GAGACTTGGCCCGACAATAA | CTCGCCGTAACATTCACCTT |
| *ZAT12* | *AT5G59820* | CATCACAACTACTATCACACCAAACTC | ATCCACCGTCGACTTGATCT |
| *nLUC* | *NA* | CCAGAACCTCGGAGTTAGCG | CTGTCCCATCTGATCTCCGC |

**Table S5. List of primers for ChIP-qPCR**

| Gene | AGI Code | Forward Primer | Reverse Primer |
|---|---|---|---|
| *UBQ10* | *AT4G05320* | TCCCTCCCTTTAAGCACCAG | TCCGGTCCTAGATCATCAGTTCA |
| *EIF4AI* | *AT3G13920* | TGTTTTGCTTCGTTTCAAGGA | GCATTTTCCCGATTACAAC |
| *ADH1* | *AT1G77120* | GCAAAACCAAATACGCCCC | TAATCTGTCCGGTGGTAGAC |
| *LBD41* | *AT3G02550* | GAGAGAGTCACAAAGATCCGCCC | GAAGAACTGGGGCCCACACTTAG |
| *HB1* | *AT2G16060* | CCATGTGCTCTGTACTGGTAATGGA | TTATACCACTTGGTGTGGTTGGC |
| *HRA1* | *AT3G10040* | GCAGTGGTTTTGGGAGCCGT | TTTGCCAAAACCAGCCCCTTG |
| *ZAT12* | *AT5G59820* | TACGCGGTGTCGCAAATCGT | TGGGTAAGGAAGTGGCAGCG |
| *ATH8* | *AT1G69880* | AGGTTTACATGCAACTTTCCGCT | CTGGTGGACCAAGTAGCCGT |
| *GSTU24* | *AT1G17170* | TCAAGTGCGCCAAAAGGAAAGA | AGGGGTTTTGAATCGCATTTTGCT |
